# Glycosylphosphatidylinositol-anchoring is required for the proper transport and glycosylation of classical arabinogalactan protein precursor in tobacco BY-2 cells

**DOI:** 10.1101/2020.10.19.346049

**Authors:** Daiki Nagasato, Yuto Sugita, Yuhei Tsuno, Rutsuko Tanaka, Maki Fukuda, Ken Matsuoka

**Affiliations:** Graduate School of Bioresource and Bioenvironmental Science, Kyushu University, Fukuoka, Japan; Chikushigaoka High School, Fukuoka, Japan; School of Agriculture, Kyushu University, Fukuoka, Japan; Faculty of Agriculture, Kyushu University, Fukuoka, Japan

**Keywords:** AGP, arabinogalactan protein, GPI anchoring, transport pathway, glycan structure

## Abstract

Arabinogalactan proteins (AGPs) are extracellular proteoglycans with many O-linked glycan chains. Precursors to many AGPs contain a C-terminal signal for the addition of a glycosylphosphatidylinositol (GPI)-anchor, but the role of this modification has not been elucidated. NtAGP1, a tobacco precursor to AGP, comprises a signal peptide, an AGP-coding region, and a GPI-anchoring signal, and it is classified as a member of the classical AGP family. Using green fluorescent protein (GFP) and sweet potato sporamin (SPO) as tags and tobacco BY-2 cells as the host, we analysed the transport and modification of NtAGP1. The fusion protein of GFP or SPO and NtAGP1 expressed in BY-2 cells migrated as a large smear on SDS-polyacrylamide gel. A confocal microscopic analysis indicated that the GFP and NtAGP1 fusion protein localized to the plasma membrane (PM) and intracellular structures. Fractionation studies of microsomes indicated that most of the fusion protein of SPO and NtAGP1 (SPO-AGP) localized to the PM. In contrast, the expression of mutants without a GPI-anchoring signal yielded several forms. The largest forms migrating as large smears on the gel were secreted into the culture medium, whereas other forms were recovered in the endomembrane organelles. A comparison of the glycan structures of the microsomal SPO-AGP and the secreted mutant SPO-AGP without a GPI-anchoring signal using antibodies against AGP glycan epitopes indicated that the glycan structures of these proteins differ. These observations indicate that a GPI-anchoring signal is required for both the proper transport and glycosylation of the AGP precursor.

## Introduction

Arabinogalactan proteins (AGPs) are plant cell wall proteoglycans with diverse functions (Pereira et al. 2015; Hromadová et al. 2021). Most of them contain many arabinogalactan (AG)-type glycan chains attached to Hyp residues that are post-translationally generated by the action of prolyl hydroxylases (Showalter et al. 2010). These proteins are classified into at least five groups based on their domain composition (Showalter et al. 2010). One of these groups consists of precursors to the classical AGPs; each of these precursors is made up of a signal peptide for the translocation of the endoplasmic reticulum (ER) membrane, glycosylation domains with many proline residues, and a signal sequence for the attachment of a glycosylphosphatidylinositol (GPI) anchor (Showalter et al. 2010; Yeats et al. 2018). Some members of other classes of AGP precursor do not contain a GPI-anchoring signal (Showalter et al. 2010).

GPI anchors are a class of lipid anchors that retain proteins at the surface of the plasma membrane (PM), especially within the sphingolipid-sterol-rich lipid domain (Liu and Fujita 2020). Based on analyses of mammalian cells and baker’s yeast, the biosynthesis of GPI anchors and their subsequent modification during transport to the PM proceed as follows. First, the biosynthesis of the GPI anchor starts at the ER, and then the pre-assembled GPI anchor is transferred from the ER to the carboxy-terminus of proteins that contain the GPI-anchoring signal to yield a GPI-anchored protein. With the assistance of p24 proteins, GPI-anchored proteins are efficiently sorted into coat protein complex II (COPII) vesicles at the ER exit site and transported to the Golgi apparatus (Bernat-Silvestre et al. 2020; Muñiz and Riezman 2016). During the subsequent transport to the PM by passing through the Golgi apparatus, the lipid part of the anchor undergoes remodeling and changes its structure (Liu and Fujita, 2020).

Most of the orthologs of genes that are indispensable for the biosynthesis and remodeling of GPI anchors in mammals and yeast are found in higher plants (Beihammer et al. 2020). Only a limited number of genes that are predicted to be involved in this step have been characterized to date (Bernat-Silvestre et al. 2021; Lin et al. 2022; Xu et al. 2022), in part because some of their null mutants are embryo-lethal (Yeats et al. 2018).

The biosynthesis of AG glycan on an AGP starts at the ER and *cis*-side of the Golgi apparatus, where Pro residues are converted to Hyp residues by the action of prolyl-hydroxylases localized in the ER and the *cis*-side of the Golgi apparatus (Yuasa et al. 2005; Velasquez et al. 2015; Parsons et al. 2018). The subsequent addition of galactose to the Hyp residue by Hyp galactosyltransferases (Oka et al. 2010; Basu et al. 2013, 2015; Ogawa-Ohnishi and Matsubayashi 2015) and the building of the AG glycan on the galactose residue in the Golgi apparatus allows the maturation of the AGP precursor (reviewed in Showalter and Basu 2106). During or after maturation, glycosylated and GPI-anchored AGPs are transported to the PM, and some of them are released to the extracellular space by the action of phospholipase in the cell wall. Several enzymes involved in AGP glycan synthesis have been characterized (reviewed in Showalter and Basu 2016), but such enzymes have not yet been characterized completely based on the structure of the AGP glycans from tobacco BY-2 cells and Arabidopsis (Tan et al. 2004; Tryfona et al. 2015; Showalter and Basu, 2016).

As both the remodeling of the GPI anchors and the maturation of AGP glycans takes place in the Golgi apparatus, there may be a linkage between these two modifications. The Golgi apparatus is an organelle located in the endomembrane transport system, and this system directs proteins to several destinations including the PM, extracellular space, and vacuoles. As such, GPI anchoring might play a role in the proper transport of proteins. However, this possibility has not been adequately investigated, especially with respect to the transport of AGP. It has been reported that GPI anchoring is indispensable for PM localisation of the Citrin-fused FLA4 protein, a fasciclin-domain containing AGP in Arabidopsis (Xue et al. 2017). In this case, most of the green fluorescence of the mutant Citrin-FLA4 lacking a GPI-anchoring signal appeared to be retained in the ER, and only a part of the green fluorescence from this protein seemed to be secreted to the extracellular space, although this partial secretion was sufficient to complement the mutant phenotype of fla4 (Xue et al. 2017). Although this observation indicates that GPI anchoring is required for the proper localisation of this protein, this observation may not be applicable to other classes of AGP; e.g., classical AGP, which does not contain a fasciclin domain, and almost all of the mature polypeptide region is predicted to be heavily glycosylated with AG glycans (Showalter et al. 2010).

Here we investigated the role of GPI anchoring in the transport and glycosylation of tobacco NtAGP1 (GenBank no. BAU61512), which is a classical AGP expressed in tobacco BY-2 cells, by expressing two different protein-tagged versions in tobacco BY-2 cells. We present evidence that GPI anchoring is required for both the proper transport and the proper glycosylation of this protein.

## Results

### Expression of the GFP-NtAGP1 fusion protein and its mutant without a GPI-anchoring signal in tobacco BY2-cells

The NtAGP1 was used as a model protein for classical AGP in this study. The amino acid sequence of this protein is nearly identical to that of *Nicotiana alata*-style AGP (Du et al. 1994). NtAGP1 is encoded by the expressed sequence tag (EST) clone BY28237 isolated from the mixture of cDNA prepared from various hormone-treated BY-2 cells (Galis et al. 2006). Four EST clones that have identical or nearly identical nucleotide sequences have been identified in our EST collections (Suppl. Table S1). Among them, three were obtained from the cDNA library from the hormone-treated BY-2 cells (Galis et al. 2006), and the fourth was from the mixture of cDNA prepared from lag-, log- and stationary growth-phase cells (Matsuoka et al. 2004). An expression analysis using microarrays (Matsuoka et al. 2004) indicated that the expression level of all of these clones was higher than average in both lag and log phases, with nearly equal levels of expression at the stationary phase (Suppl. Table S1). A BLASTN search against tobacco EST sequences using the nucleotide sequence for NtAGP1 identified 43 EST sequences (Suppl. Table S2). These sequences were obtained from a number of different cDNA libraries from various tobacco organs, including a female gamete, two-celled proembryo, leaf, flower, root, germinating seeds, and mixture of several tissues. NtAGP1 and its orthologous genes are thus expressed in many tobacco organs and at major growth phases of tobacco BY-2 cells.

The precursor to NtAGP1 consists of 132 amino acids (aa): a 21-aa signal peptide, an 86-aa glycosylation domain, and a 25-aa GPI-anchoring signal. Within the glycosylation region there are 23 Pro residues, all of which are surrounded with amino acids that make up the targeting sites for prolyl hydroxylases (Shimizu et al. 2005). Among these proline residues, 15 fit to the arabinogalactosylation site (Shimizu et al. 2005), and the other eight are contiguous prolines that might form the site for the attachment of oligo-arabinose (Xu et al. 2008). Because most of the distances between the proline residues in NtAGP1 are ≤5 aa, and because the minimum length of a peptide epitope is six aa residues (Singh et al. 2013), we thought that it would be almost impossible to generate an antibody against the peptide sequence of NtAGP1 to detect the native protein in tobacco tissues and cells. We thus adopted the alternative approach of expressing fusion proteins consisting of NtAGP1 and protein tags and characterising these proteins in tobacco BY-2 cells.

We first generated a construct for the expression of a fusion protein containing green fluorescence protein (GFP) between the signal peptide and the subsequent region of NtAGP1 (GFP-AGP). We also generated a mutant of this protein without a GPI-anchoring signal (GFP-AGPΔC) and a signal peptide with GFP only (GFP) (Fig. 1A). These proteins as well as the fusion proteins of the PM water channel and GFP (PIP-GFP: Yamauchi et al. 2003) were expressed in tobacco BY-2 cells, and the resulting transformed cells were cultured in suspension.

**Fig. 1.**
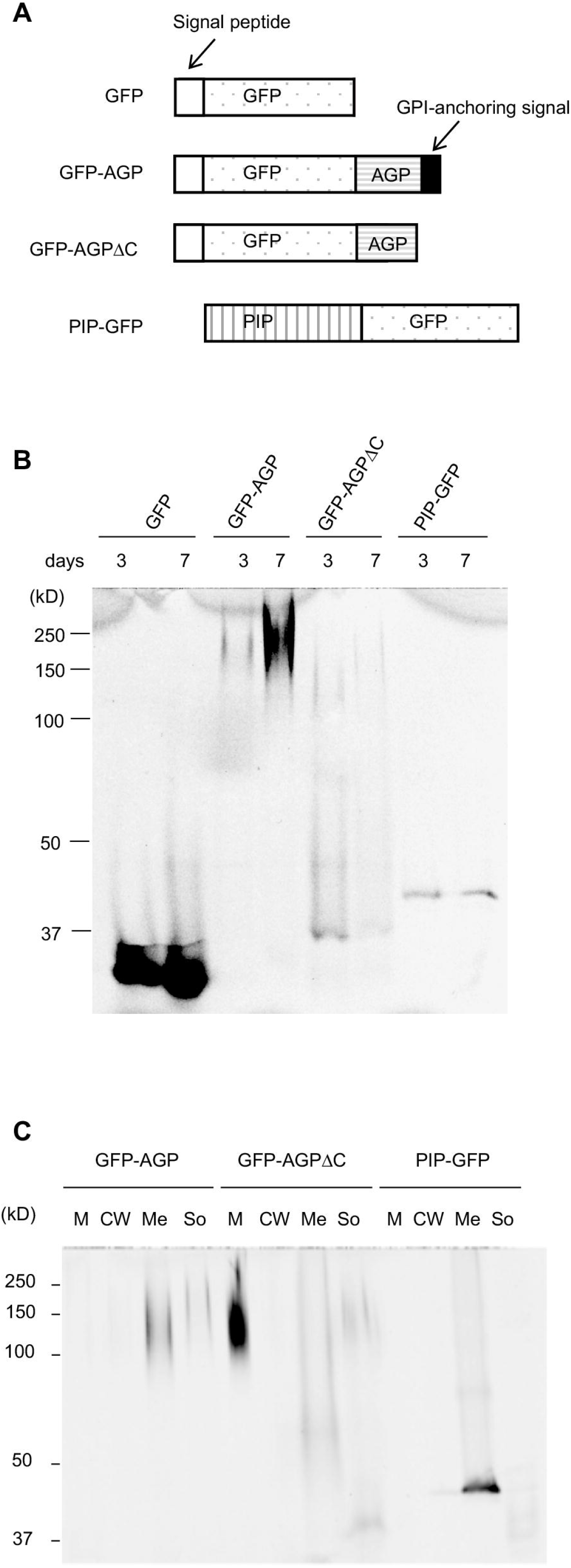
Expression of the GFP-AGP fusion protein and its mutant without a GPI-anchoring signal in tobacco BY-2 cells. **A:** Schematic representation of the GFP fusion constructs. **B:** Accumulation of GFP fusion proteins in the culture of transformed tobacco cells. Total cell lysates were prepared from cells from 3-day and 7-day cultures, then separated by SDS-PAGE, and the GFP fluorescence in the gel was recorded. Each lane contained 3.6 μg of protein. **C:** Fractionation of 7-day cultures and the distribution of the GFP fusion proteins. Each lane contained proteins corresponding to 40 μL of cell culture. M: medium fraction, CW: cell wall fraction, Me: membrane fraction from total cell lysate, So: soluble fraction from total cell lysate.

We then analysed the GFP fluorescence of proteins in these transformed cell cultures that were grown for either 3 days (to the logarithmic growth phase) or 7 days (to the stationary growth phase) after the separation of proteins by sodium dodecyl sulfate-polyacrylamide gel electrophoresis (SDS-PAGE) and fluorescence recording (Fig. 1B). The fluorescence of GFP-AGP migrated as a large smear at around the 200-kDa position, and the intensity of this signal was stronger at 7 days. In the case of GFP-AGPΔC, at least three smear bands were observed: two bands that migrated to the 120-kDa and 48-kDa positions, and a relatively sharp band that migrated to around the 37-kDa position, the size of which corresponded to the non-glycosylated form of this approx. 35-kDa protein (Fig. 1B). Among these proteins, the largest form migrated more slowly at 7 days than that at 3 days on the SDS-polyacrylamide gel. The control proteins, GFP and PIP-GFP, migrated to the corresponding position on the SDS-polyacrylamide gel, as expected.

To determine the localisation of these GFP fusion proteins, cultures of the transformed cells were separated into medium, cell wall, cellular membrane, and cellular soluble fractions. Proteins in these fractions were separated, and the distribution of the GFP fusion proteins was analysed (Fig. 1C). GFP-AGP was recovered predominantly in the membrane fraction and to a lesser extent in the soluble fraction, and trace amounts were present in the cell walls and culture medium. In the case of GFP-AGPΔC, most of the large smear was recovered in the medium, and a small amount of this form was also found in the membrane and soluble fractions. In contrast, almost all of the 48-kDa form was recovered in the membrane fraction, and the smallest 37-kDa form was recovered in the soluble fraction. In the case of PIP-GFP, which was used as the fractionation control, most of this protein was recovered in the membrane fraction with a trace amount in the cell walls, and faint signals of smaller forms, possibly truncated ones, were detected in the soluble fraction.

We next addressed the distribution of the GFP fluorescence from 7-day-old cells expressing either GFP-AGP or GFP-AGPΔC by a confocal laser scanning microscopy (CLSM) examination (Fig. 2). Almost all of the green fluorescence in the GFP-AGP-expressing cells was localized at the cell surface, possibly at the PM, and a weak signal was observed in the vacuoles as well as in unidentified punctate structures (Fig. 2A, GFP-AGP). The intensities of fluorescence varied in the cells, and the cell-surface signal seemed nonuniform. In contrast to the cell-surface pattern of GF-AGP signal, cells expressing GFP-AGPΔC did not show such a pattern of fluorescence; they did exhibit weak green fluorescence in the vacuoles with an unidentified punctate structure with green fluorescence close to the cell wall (Fig. 2, GFP-AGPΔC).

**Fig. 2.**
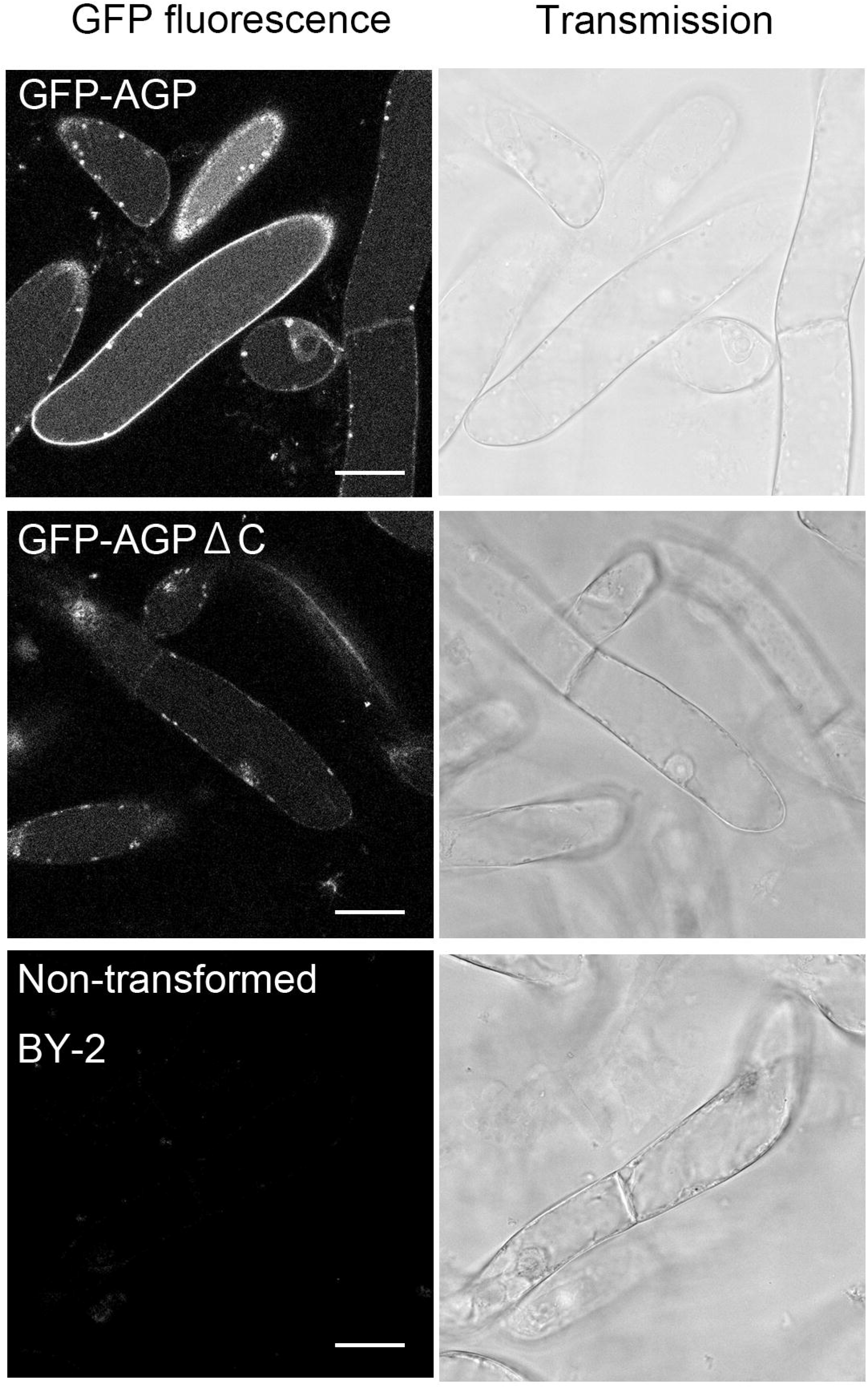
Confocal localisation study of GFP-AGP and GFP-AGPΔC in transformed tobacco BY-2 cells. Confocal image of 7-day-old transformed tobacco cells expressing GFP-AGP, GFP-AGPΔC, or non-transformed cells. *Left:* Confocal fluorescence image and (*right*) the corresponding DIC image. Bars: 30 μm.

To determine whether the GFP-AGP signal was localized in the PM, we incubated the cells for 5 min with FM 4-64 to visualize the plasma membrane, and both the GFP and FM 4-64 fluorescence were recorded (Fig. 3A). The GFP and FM 4-64 fluorescence colocalized well at the edges of the cells. The quantification of signal intensities on a line along the short axis at the middle of the cell (Fig. 3A, merged image, yellow line) indicated that the majority of the GFP fluorescence merged with the FM 4-64 fluorescence (Fig. 3A graph), although some GFP signal was also observed at the inside of the FM 4-64 signal.

**Fig. 3.**
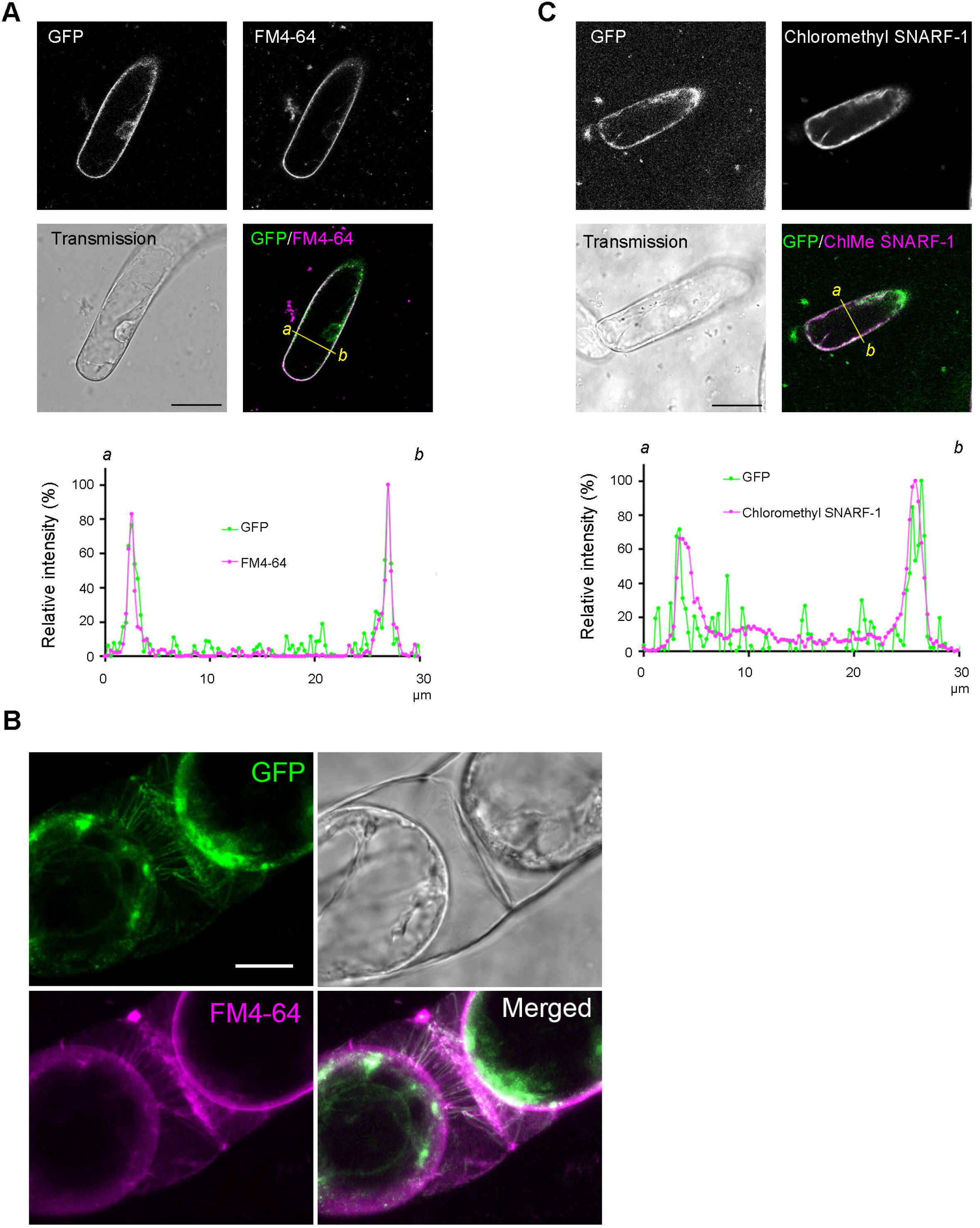
Comparison of the localisation of GFP-AGP and plasma membrane or cytosol markers. **A:** Comparison of GFP-AGP and plasma membrane marked with FM4-64. Cells expressing GFP-AGP were incubated for 5 min with FM 4-64, then the fluorescence of both GFP and FM 4-64 was recorded using CFLM. The intensities of the fluorescence signals along the *yellow line* in the merged image were quantified. Normalized (maximum as 100%) intensities are shown in the graph. **B:** Confocal image of cells expressing GFP-AGP after staining with FM4-64 and plasmolysis. Composite images from 23 Z-stack images encompassing 7.96 μm are shown for GFP-AGP and FM 4-64 signals. **C:** Comparison of GFP-AGP and cytoplasm marked with 5- (and 6-) chloromethyl SNARF-1. Cells expressing GFP-AGP were incubated with 5- (and 6-) chloromethyl SNARF-1 acetate (SNARF) for 5 min; confocal images were collected, and the intensities of the signals were quantified as in panel A. Bars: 30 μm in A and C, and 10 μm in B.

To confirm whether a significant proportion of green fluorescence from GFP-AGP-expressing cells was localized to the PM, we stained the cells with FM 4-64 and observed the cells under plasmolysis (Fig. 2B). The clear lines of a green fluorescence signal, which corresponded to Hechtian strands, were visible between the PM and cell walls. At the same position, Hechtian strands with FM 4-64 fluorescence were present. Some of the fluorescence also seems attached to the cell wall, and some was present in the cell. These observations suggested that the majority of GFP-AGP was localized to the PM although some of it was present in intracellular structures.

To further confirm whether the majority of the GFP-AGP signal at the edge of the cells indicates the PM localisation, we stained the cytoplasm with 5- (and 6-) chloromethyl SNARF-1 acetate (SNARF) and compared the localisation of the GFP fluorescence with that of SNARF (Fig. 3C). The majority of the SNARF fluorescence localized close to the cell wall, in a pattern which was similar to that of GFP. However, a close inspection of the merged images and the quantification of signal intensities on a line along the short axis at the middle of the cell (Fig. 3B, merged image, yellow line) indicated that the peaks of the GFP fluorescence were one or two pixels outward from the peak of SNARF fluorescence (Fig. 3B graph). These observations further confirmed that a significant proportion of GFP-AGP was localized to the PM.

As described above, optical section images suggested that the GFP fluorescence of GFP-AGP on the plasma membrane was not distributed evenly on the plasma membrane. To assess the non-uniform localisation of GFP-AGP at the PM, we collected Z-stack images by CLSM and then reconstituted a 3D movie (Suppl. Movie S1). The nonuniform distribution of GFP fluorescence was apparent in the reconstituted 3D movie. These data indicated that GFP-AGP was predominantly localized to the PM, with a nonuniform distribution.

Although these data suggested that the majority of the NtAGP1 is localized to the PM, we could not confirm this localisation based on the result of the expression of only one fusion protein, and it was difficult to further characterize GFP-AGP, for several reasons. (1) The green fluorescence from GFP-AGP was unstable in the stably transformed tobacco BY-2 cells, and within six weeks of subculture, almost all of the cells stopped emitting green fluorescence. (2) The signal peptide-GFP fusion protein was not fully secreted to the culture medium from tobacco BY-2 cells (Mitsuhashi et al. 2000). Thus, GFP alone may not be a good cargo for secretion. (3) We have a fluorescence pulse-chase system that uses a photo-convertible fluorescence protein, mKikGR (Habuchi et al. 2008) which has a higher-order structure that is similar to that of GFP, to monitor the turnover of proteins (Abiodun and Matsuoka 2013). We thus attempted to apply this system to address the transport of NtAGP1 by expressing the fusion protein of mKikGR and NtAGP1 in tobacco BY-2 cells. Unfortunately, the fusion protein migrated with three distinct bands: a weak large smear that migrated at a position similar to that of GFP-AGP, a predominant band around 60 kDa, and a weaker band around 50 kDa (Suppl. Fig. S1). We therefore tested the transport and modification of NtAGP1 using a protein that is structurally distinct from GFP as a tag in order to determine the localisation and transport of this protein and its mutant lacking a GPI-anchoring signal.

### Expression and localisation of the sporamin-NtAGP1 fusion protein and its mutant lacking an GPI-anchoring signal in tobacco BY-2 cells

We used sweet potato sporamin (SPO) as another protein tag to address the transport, localisation, and modification of NtAGP1 and its derivative lacking a GPI-anchoring signal. Sporamin is a monomeric soluble and non-glycosylated storage protein of sweet potato (Maeshima et al. 1985) localized in the vacuoles (Hattori et al. 1988). The expression of a mutant precursor to sporamin, which does not contain the propeptide region but does contain the signal peptide for secretion, allowed the efficient secretion of this protein to the medium from transformed tobacco BY-2 cells (>90% of this pulse-labeled protein is secreted within 2 h of chase) (Matsuoka and Nakamura 1991). The junction region between the propeptide and the mature sporamin surrounding the 36th Pro residue consists of a cryptic Hyp O-glycosylation site for AG, and thus both a wild-type precursor to sporamin as well as mutant precursors with a disrupted vacuolar targeting signal are O-glycosylated when expressed in tobacco BY-2 cells (Matsuoka et al. 1995, Matsuoka and Nakamura 1999).

The efficiency of the secretion of mutants with a disrupted vacuolar targeting signal is comparable to that of the mutant without the propeptide (Matsuoka and Nakamura 1999). In other words, the secretion efficiency of sporamin is not affected by the presence or absence of O-glycosylation. We thus chose sporamin as another protein tag to analyse the transport and modification of NtAGP1 and its mutant lacking a GPI-anchoring signal.

The expression constructs for the signal peptide-sporamin-mature NtAGP1 with a GPI-anchoring signal (SPO-AGP) were expressed in tobacco BY-2 cells, as were the expression constructs for the signal peptide-sporamin-mature NtAGP1 without a GPI-anchoring signal (SPO-AGPΔC) (Fig. 4A). Cell and medium fractions were prepared from stationary-grown transformed BY-2 culture, and then the intra- and extra-cellular localisations of sporamin or its fusion proteins in these fractions were examined by immunoblotting either directly without immunoprecipitation (Fig. 4B) or after target proteins were recovered by immunoprecipitation (Fig. 4C). Immunoblotting without immunoprecipitation gave cross-reacted protein bands in proteins from non-transformed cells (Fig. 4B, WT). Such signals were absent after immunoprecipitation (Fig. 4C, WT). However, our comparison of the intensities of sporamin-related signals on many immunoblots obtained under both these conditions indicated that the recovery by immunoprecipitation varied among the proteins, especially SPO-AGP. We thus did not use immunoprecipitation for the subsequent quantitative analysis, and we always included a negative control from non-transformed cells.

**Fig. 4.**
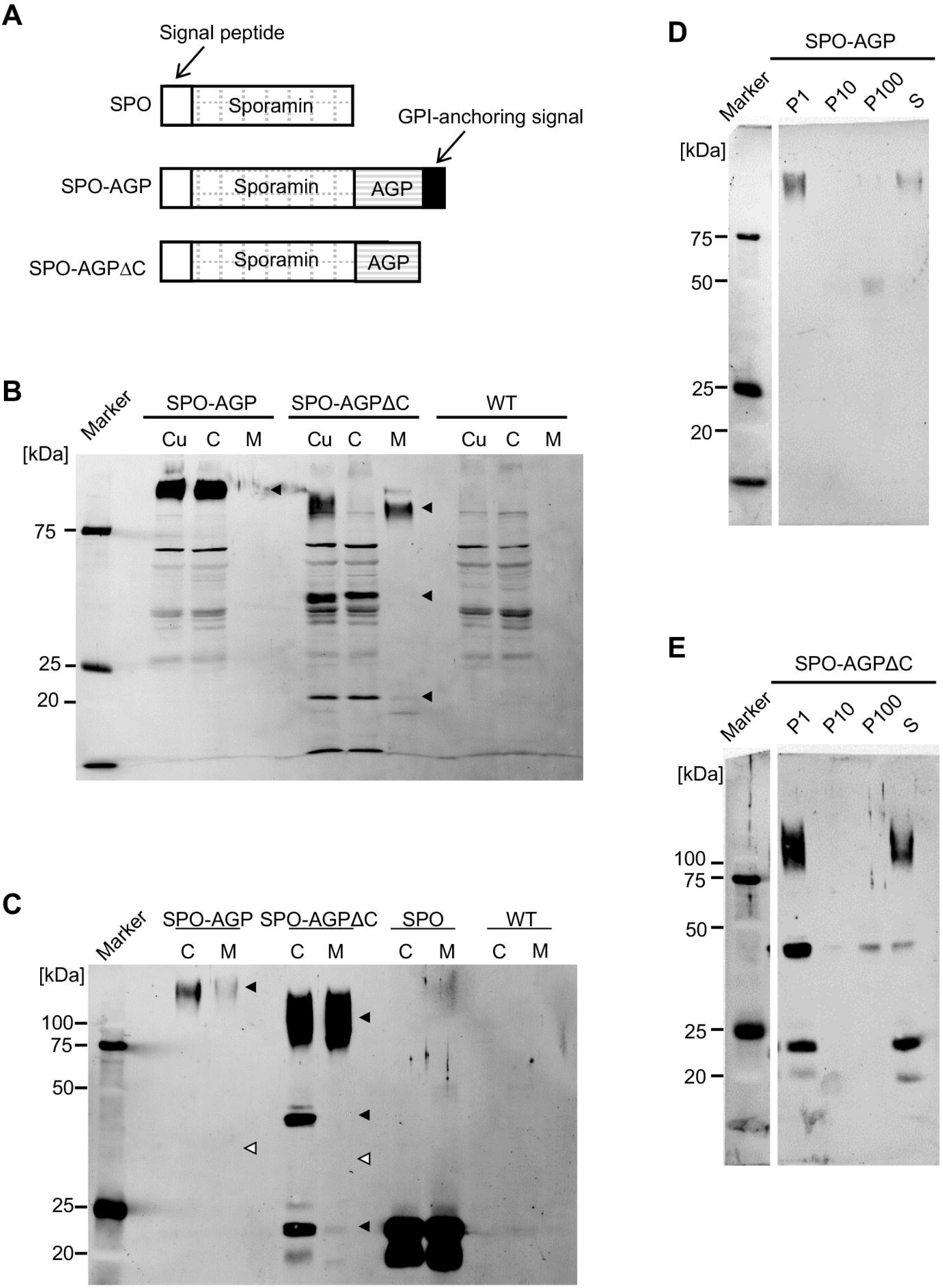
The expression, secretion, and subcellular localisation of the sporamin-AGP fusion protein and its mutant lacking a GPI-anchoring signal in tobacco BY-2 cells. **A:** Schematic representation of the sporamin fusion constructs. **B:** Expression and secretion of SPO-AGP and SPO-AGPΔC. Seven-day-old cultures (Cu) of transformed BY-2 cells expressing SPO-AGP and SPO-AGPΔC and non-transformed BY-2 cells (WT) were separated into cells (C) and medium (M). The proteins in these fractions, corresponding to 10 μL of cell culture, were separated by SDS-PAGE, and sporamin-related polypeptides were detected by using an anti-SDS-denatured sporamin antibody. *Closed arrowheads:* transgene-dependent polypeptides. **C:** Detection of sporamin-related polypeptides after immunoprecipitation. Sporamin-related polypeptides were recovered by immunoprecipitation using immobilized anti-native sporamin, and the recovered proteins were analysed by immunoblotting as described in the Materials and Methods section. Each lane contained sporamin-related polypeptides from 0.5 mL of 7-day-old culture. *Closed arrowheads:* the migration position of transgene-dependent polypeptides detected in panel B. *Open arrowheads:* the calculated migration position of non-glycosylated polypeptides. **D:** Subcellular fractionation study of SPO-AGP-expressing cells. Fractionation of cells was carried out by differential centrifugation as described in the Materials and Methods, and sporamin-related polypeptides in the fractions were recovered by immunoprecipitation, separated by SDS-PAGE, and detected by immunoblotting. P1: the 1,000*g* precipitate containing cell walls and unbroken cells, P10: the 10,000*g* precipitate containing most of the mitochondria, P100: the 100,000*g* precipitate containing microsomes, S: the 100,000*g* supernatant containing soluble proteins in the cytoplasm, vacuoles and periplasmic space. **E:** Subcellular fractionation study of SPO-AGPΔC-expressing cells. The subcellular fractionation and detection of sporamin-related polypeptides were carried out as in panel D.

SPO-AGP was detected as a smear band close to the top of the SDS-polyacrylamide gel (Fig. 4B, C). This observation indicated that SPO-AGP was glycosylated. Most of the smear-migrating protein was recovered in the cell fraction (Fig. 4B). In contrast, SPO-AGPΔC was detected as three distinct species with different migration positions; a large form that migrated at the 90–120-kDa position, an intermediate form at the 36-kDa position and a small one at the 22-kDa position (Fig. 4B, C). A specific band that migrated at the front of the electrophoresis gel may have been the degradation product of SPO (Fig. 4B). A large form of SPO-AGPΔC was detected in both the cell and medium fractions. Because the calculated molecular weight of non-glycosylated SPO-AGPΔC is 27.5, this observation indicated that most of the large-form SPO-AGPΔC was glycosylated and secreted into the extracellular space. The intermediate and small forms of SPO-AGPΔC were detected in cell fractions. This observation suggests that they were localized in the cells. The small form may be the degradation product, since its size (22 kDa) was similar to that of SPO (Fig. 4C) and smaller than the calculated size of the non-glycosylated form of SPO-AGPΔC.

To reveal the intracellular localisation of SPO-AGP and SPO-AGPΔC, we performed subcellular fractionation by differential centrifugation. Cell lysates were centrifuged at 1,000*g* and the precipitate, defined as the P1 fraction (which contained unbroken cells, nuclei, cell walls and cell wall-attached PM) was collected. The supernatant was centrifuged at 10,000*g* and the precipitate (P10), which contained most of the mitochondria and plastids, was collected. The resulting supernatant was further centrifuged at 100,000*g* and the obtained precipitate (P100) contained microsomal membranes. The supernatant, which was defined as the S fraction, was also collected. SPO-AGP was detected in the P1, P100, and S fractions (Fig. 4D). In the case of SPO-AGPΔC, the large and small forms were detected in the P1 and S fractions, while the intermediate form was observed in all fractions (Fig. 4E).

To address the localisation of SPO-AGP, which was recovered in the P100 fraction, we prepared microsomal membranes from transformed BY-2 cells expressing SPO-AGP using buffers containing either MgCl_2_ or EDTA, and we then separated the membranes by equilibrium sucrose density gradient centrifugation in the presence or absence of Mg^2+^, respectively. The distribution of SPO-AGP in these separated fractions was examined by immunoblotting (Fig. 5A). The distributions of marker proteins were also analysed by immunoblotting with specific antibodies: Sec61 for the ER (Yuasa et al. 2005), vacuolar pyrophosphatase (V-PPase) for tonoplast (Toyooka et al. 2009), GLMT1 for the Golgi apparatus (Liu et al. 2015), plasma membranous ATPase (P-ATPase) for the PM (Toyooka et al. 2009) and the PM water channel (PIP) for the PM (Suga et al. 2001). In the presence of Mg^2+^ (Fig. 5A, +MgCl_2_), SPO-AGP migrated in several fractions, mainly in the fractions corresponding to 41–44(w/w)% sucrose. Quantification of the intensities of these immunoblot signals indicated that the migration pattern of SPO-AGP was similar to those of P-ATPase and PIP, the markers of the PM. In the absence of Mg^2+^ (Fig. 5A, +EDTA), SPO-AGP also migrated with a peak at around 41–43(w/w)% sucrose. This migration pattern was also similar to those of P-ATPase and PIP. These fractionation data indicated that SPO-AGP was localized to the PM.

**Fig. 5.**
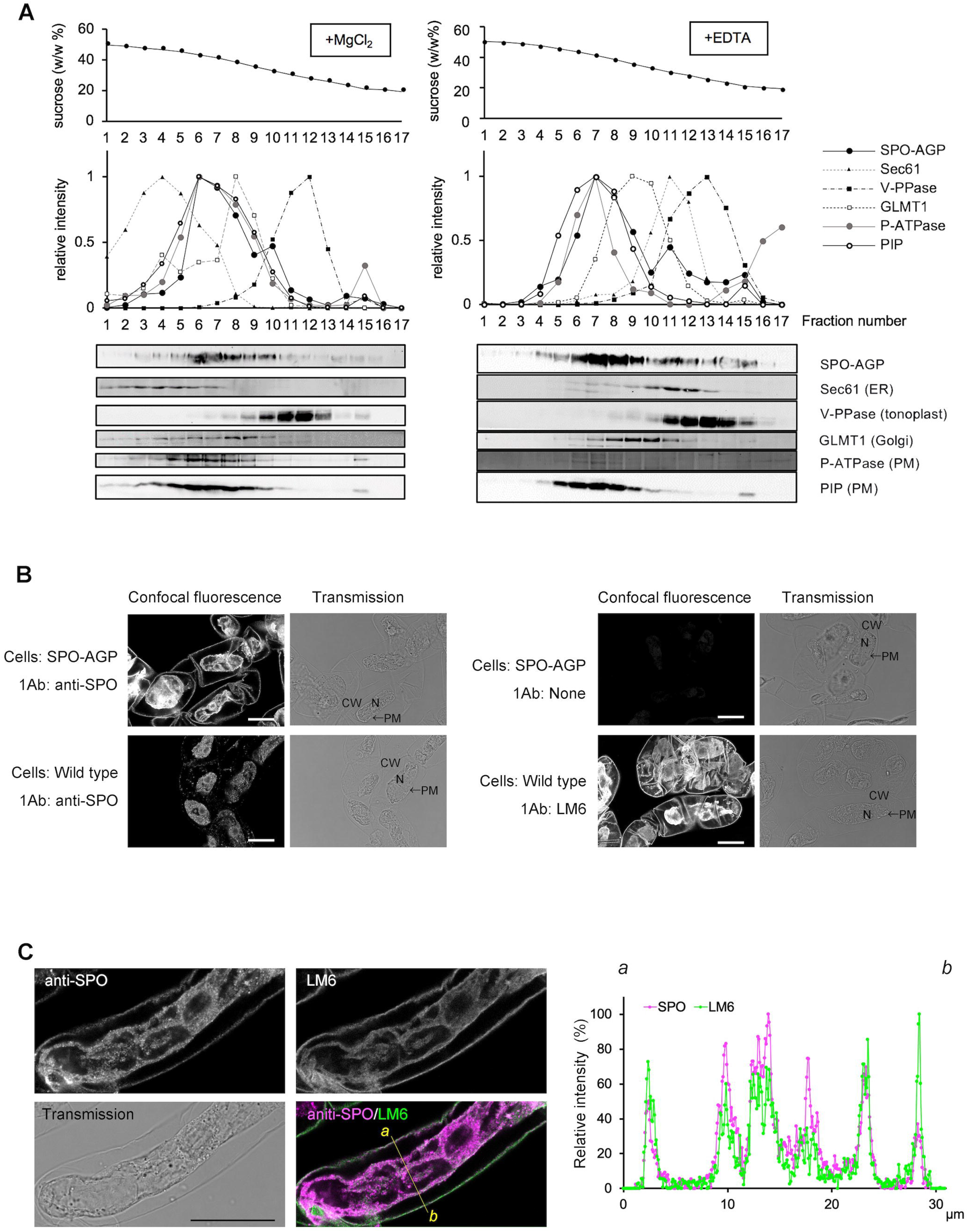
Intracellular distribution of SPO-AGP. **A:** Microsomes were prepared from SPO-AGP-expressing cells in the presence of Mg^2+^ or EDTA and subjected to isopycnic sucrose density gradient ultracentrifugation. The resulting gradients were fractionated from the bottom into 17 fractions. The concentration of sucrose in the gradient is shown at the top. The distribution of marker proteins was analysed by immunoblotting with specific antibodies: P-ATPase and PIP for PM, Sec61 for the ER, GLMT1 for the Golgi apparatus, and V-PPase for the vacuolar membrane. The distribution of SPO-AGP, which migrated close to the top of the gel, was analysed by immunoblotting using an antibody against sporamin. *Middle panels:* the relative distribution of SPO-AGP and marker proteins after quantification of the signals on blots. *Bottom panels:* the immunoblot results. **B:** Immunostaining of SPO-AGP-expressing or non-transformed cells with anti-native SPO antibody or anti-AGP glycan LM6. C: Colocalisation of SPO-AGP and anti-AGP glycan epitope. SPO-AGP-expressing cells were stained with both anti-native SPO antibody and anti-AGP glycan LM6 with appropriate secondary antibodies. *Left:* Confocal images of both stained and corresponding transmission images. Intensities of the fluorescence signals along the *yellow line* in the merged images were quantified. Normalized (maximum as 100%) intensities are shown in the graph.

To confirm the PM localisation of SPO-AGP with a different method and to compare the localisation of this protein with endogenous AGP, we fixed transformed cells expressing SPO-AGP with formaldehyde and stained them with anti-sporamin or LM6 (Willats et al. 1998), which is a monoclonal antibody against oligo-arabinose epitope in both AGP glycan and rhamnogalacturonan I in pectin (Fig. 5B). LM6 recognized the glycan on SPO-AGP by immunoblotting (see below). For the control, non-transformed BY-2 cells were fixed and stained as well (Fig. 5B). During the fixation most of the cells shrank and separated from the cell walls, and thus the localisation at the cell wall and cells was easily distinguished, although some of the plasma membrane proteins were cross-linked to cell walls during fixation.

The staining of SPO-AGP-expressing cells with anti-native SPO antibody showed a clear signal at the cell wall, PM, and punctate structures in the cell, although a weak intracellular puncta signal was also observed when non-transformed cells were stained with this antibody (Fig. 5B). The pattern of staining of the SPO-AGP-expressing cells with anti-sporamin antibody resembled the pattern of staining of wild-type cells with LM6 (Fig. 5B), although the SPO-AGP signal at the cell wall showed a more uneven distribution compared to that of LM6. Specificities of the staining including SPO-AGP dependency were confirmed by staining both SPO-AGP-expressing cells and wild-type cells with either and both antibodies (Suppl. Figs. S2, S3). The double immunostaining of SPO-AGP-expressing cells with anti-native SPO antibody and LM6 (Fig. 5C) revealed that both signals were colocalized well, although the SPO-AGP signal in the cells was more punctate compared to that of LM6. A comparison of signal intensities on a line along the short axis at the middle of the cell (Fig. 5C, merged image, yellow line) further confirmed that the majority of the SPO-AGP signal merged with the LM6 signal, especially at the cell wall and PM (Fig. 5C graph).

To address the stability of SPO-AGP, we measured the level of SPO-AGP after protein synthesis was stopped in the presence of cycloheximide. Although the average intensity of the SPO-AGP signal under the cycloheximide condition was lower than the control value, the difference was not significant (Suppl. Fig. S4). These observations suggest that SPO-AGP is not an unstable protein and that is localized mainly at the PM.

### Intracellular localisation of SPO-AGPΔC

We prepared protoplasts from transformed BY-2 cells expressing SPO-AGPΔC in order to determine which forms of SPO-AGPΔC were localized outside of the PM (Fig. 6A). The large form of SPO-AGPΔC was detected only in cells and was not detected in protoplasts. The intermediate-form protein was detected in both cells and protoplasts, although the amount was smaller in protoplasts than in cells. The small form of SPO-AGPΔC was detected in both fractions in nearly equal amounts. This result indicated that the large-form SPO-AGPΔC was localized in the periplasm and cell wall; the intermediate SPO-AGPΔC was localized in both the periplasm and the cells, and the small SPO-AGPΔC was localized almost exclusively in the cells.

**Fig. 6.**
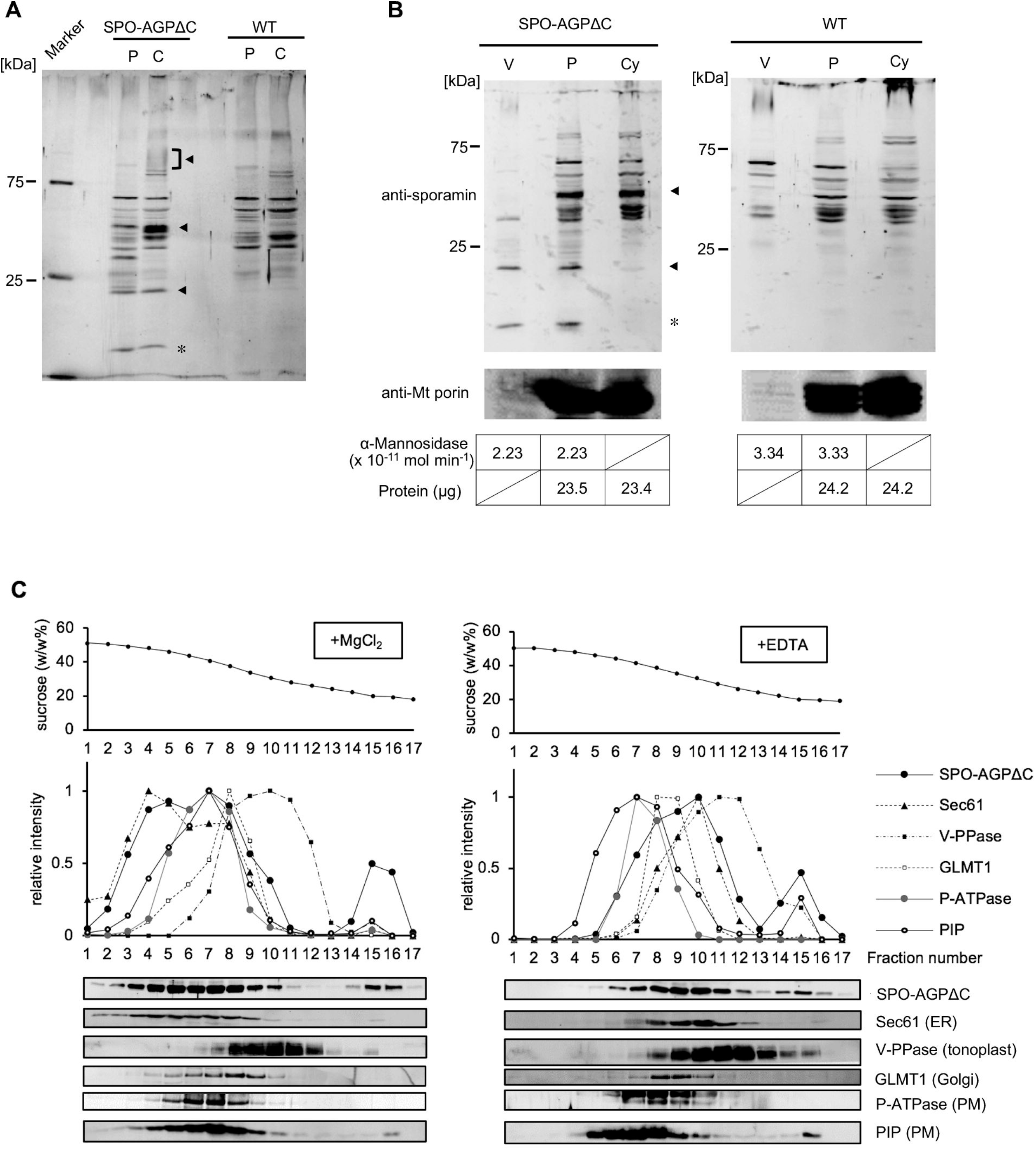
Distribution of different forms of SPO-AGPΔC in the cell. **A:** Distribution of different forms in cells and protoplasts. Protoplasts were prepared from SPO-AGPΔC-expressing cells by digesting the cell walls as described in the Materials and Methods. Proteins in cells and protoplasts were separated by SDS-PAGE, and sporamin-related polypeptides were detected by immunoblotting. As a control, protoplasts were prepared from non-transformed cells (WT), and cross-reactive polypeptides against anti-sporamin were analysed by immunoblotting. Each lane contained 25 μg of protein. *Closed arrowheads:* the migration position of the transgene-dependent polypeptides. *Asterisks:* a possible degradation product of sporamin that was occasionally observed on immunoblots. **B:** The small form of SPO-AGPΔC was localized in the vacuoles. To assess the localisation of the intermediate and small SPO-AGPΔC proteins, protoplasts were separated into vacuoplasts and cytoplasts, and the distribution of SPO-AGPΔC proteins was analysed by immunoblotting. A nearly equal amount of the activity of α-mannosidase, which is a vacuolar marker enzyme, was loaded in the vacuoplast and protoplast lanes, and nearly equal amounts of proteins were loaded into the protoplast and cytoplast lanes. As a fractionation control, mitochondrial porin was also detected by immunoblotting. **C:** The distribution of the intermediate form of SPO-AGPΔC within endomembrane organelles. Microsomes were prepared from SPO-AGPΔC-expressing cells in the presence of Mg^2+^ (*left*) or EDTA (*right*), subjected to isopycnic sucrose density gradient ultracentrifugation, fractionated, and analysed for the distribution of intermediate SPO-AGPΔC and marker proteins as described in the Fig. 5A legend.

To address whether some of the intracellular SPO-AGPΔC was localized in vacuoles, we separated protoplasts into vacuoplast and cytoplast fractions and analysed the amounts of the different forms of SPO-AGPΔC in these fractions. The vacuoplasts are composed of vacuoles surrounded by the PM, and cytoplasts are composed of cytoplasm, nuclei, and PM. We measured the protein concentration and assessed the α-mannosidase activity in the protoplast and cytoplast fractions. We then used SDS-PAGE to separate the proteins that were present in the protoplast and cytoplast fractions in nearly equal amounts and the proteins that were present in the protoplast and vacuoplast fractions in amounts that are almost equivalent to the levels of α-mannosidase activity. The different forms of SPO-AGPΔC in these fractions were detected by immunoblotting (Fig. 6B). Mitochondrial porin was used as a fractionation control. The intermediate SPO-AGPΔC was detected in the protoplast and cytoplast fractions in nearly equal amounts relative to the protein, whereas the small form was detected in both the protoplast and vacuoplast fractions in amounts similar to the activity of α-mannosidase. This result suggests that the small SPO-AGPΔC was localized in vacuoles and the intermediate SPO-AGPΔC was localized in other intracellular compartments.

To investigate the localisation of the intermediate SPO-AGPΔC in cells, we prepared microsomal fractions from transformed BY-2 cells expressing SPO-AGPΔC in buffers containing either MgCl_2_ or EDTA. The microsomal membranes were separated by equilibrium sucrose density gradient centrifugation in the presence or absence of Mg^2+^, respectively. The distribution of the intermediate SPO-AGPΔC in these separated fractions was examined by immunoblotting and compared to the distributions of marker proteins in the secretory organelles (Fig. 6C). In the presence of Mg^2+^ (Fig. 6C, +MgCl_2_), the intermediate SPO-AGPΔC migrated in several fractions with a peak at around 39–42(w/w)% sucrose. This migration pattern was similar to that of Sec61, a marker of the ER membrane. In the absence of Mg^2+^ (Fig. 6C, +EDTA), the intermediate SPO-AGPΔC also migrated with a peak at around 31–34(w/w)% sucrose, and the peak shifted to a lower-density position compared to that in the presence of Mg^2+^. This shift corresponded to that of the migration pattern of Sec61 (an ER marker). These fractions did not contain P-ATPase or PIP but did contain GLMT1, a marker of the Golgi apparatus. These fractionation data suggest that the intermediate form of SPO-AGPΔC is localized predominantly in the ER and partly in the Golgi apparatus.

### NtAGP1 is a GPI-anchored protein

The NtAGP1 precursor contains a GPI-anchoring signal at its C-terminus. Our present analyses demonstrated that GFP-AGP and SPO-AGP are localized to the PM (Figs. 2, 5). To determine whether the GPI-anchoring signal on the NtAGP1 precursor is functional and whether the GPI anchor attached to NtAGP1 was responsible for attaching this protein to the PM, we analysed the distribution of these fusion proteins recovered in the microsomal fraction by two-phase separation using the nonionic detergent Triton X-114. This nonionic detergent can be used to make a uniform solution at low temperature, which can then be separated into two phases, such as an aqueous (aqu) phase and a detergent-rich (det) phase, at higher temperature. By taking advantage of this characteristic of the nonionic detergent, a cell extract can be separated into soluble and peripheral membrane proteins that can be recovered in the aqu phase, plus integral membrane proteins as well as lipid-anchored proteins that can be recovered in the det phase (Bordier 1981).

Approximately half of the GFP-AGP (Fig. 6A, left) as well as almost all of the SPO-AGP (Fig. 6B, top left) and almost all of the PIP-GFP (an integral membrane protein used as an experimental control; Fig. 6A, right), was recovered in the det fraction. Under the same separation condition, almost all of the smear-migrating GFP-AGPΔC and SPO-AGPΔC were recovered in the aqu fraction. These observations suggested that about half of the GFP-AGP and almost all of the SPO-AGP were localized to the PM with their GPI anchors.

To address whether this membrane-association is actually mediated by the GPI anchor attached to these proteins, we carried out two-phase separation experiments in the presence of phosphatidylinositol-specific phospholipase C (PI-PLC), which can cleave the phosphodiester bond in a GPI anchor (reviewed by Hopper 2001). In the presence of PI-PLC, almost all of the GFP-AGP in the microsomal fraction was recovered in the aqu phase (Fig. 6C, right). Likewise, about half of the SPO-AGP that was recovered in the TX-114 phase was recovered in the det phase (Fig. 6D, top). Under the same condition, vacuolar pyrophosphatase, which is an integral membrane protein used as a control, was recovered almost exclusively in the det phase (Fig. 6D, bottom). These observations indicated that (*i*) roughly half of both GFP-AGP and SPO-AGP was anchored to the PM by a GPI anchor that was sensitive to PI-PLC, and (*ii*) NtAGP1 is a GPI-anchored protein at the PM.

### GPI anchoring is required for the proper assembly of AGP glcyan

We showed that the migration positions of the smear-migrating SPO-AGP and SPO-AGPΔC on SDS-polyacrylamide gel were different (Fig. 4A, B). We considered that this difference might be due to the difference in the AG glycan chains on these proteins. To address this possibility, we examined the reactivity of anti-AG glycan monoclonal antibodies (mABs) to these proteins. Among the mABs that we tested, SPO-AGPΔC was specifically recognized by LM2 (Fig. 7B). Both proteins were recognized by LM6 (Fig. 7C). Other antibodies, namely PN16.4B4 (Norman et al. 1986), CCRC-M7 (Steffan et al. 1995), and MAC204 and MAC207 (Bradley et al. 1988), did not recognize either protein. Because LM2 recognizes terminal β-glucuronic acid (β-GlcA) (Yates et al. 1996), the absence of the recognition of SPO-AGP by this antibody indicated that such a structure was absent in the glycan in SPO-AGP. This result also suggested that GPI anchoring is required for the proper assembly of AGP glycan.

**Fig. 7.**
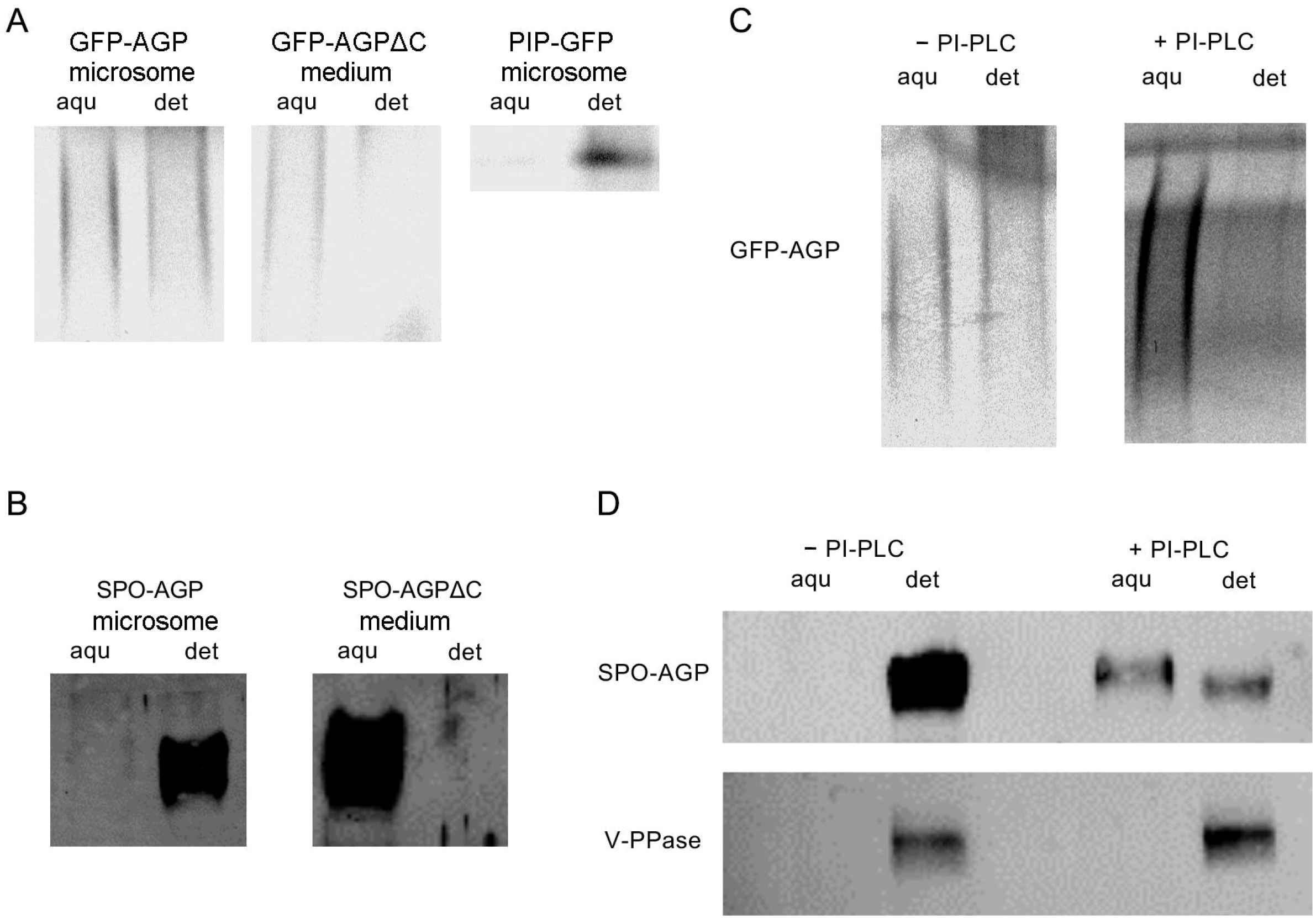
Both GFP-AGP and SPO-AGP are GPI-anchored proteins. **A:** Triton X-114 two-phase partition assay of GFP-AGP, GFP-AGPΔC and PIP-GFP. Microsomes from GFP-AGP- or PIP-GFP-expressing cells and the culture medium of GFP-AGPΔC-expressing cells was subjected to the two-phase separation assay as described in the Materials and Methods. Proteins in these two phases were separated by SDS-PAGE, and the fluorescence of GFP in the gel was recorded. Each lane contained proteins corresponding to an equal amount of microsomes or medium. aqu: aqueous phase, det: detergent phase. **B:** Triton X-114 two-phase partition assay of SPO-AGP and SPO-AGPΔC. Two-phase partition was carried out as described in the legend for panel A. Proteins were separated by SDS-PAGE and detected by immunoblotting using anti-sporamin antibody. **C:** PI-PLC digestion of GFP-AGP. The two-phase partition assay was carried out in the absence and presence of PI-PLC. **D:** Two-phase partition assay of SPO-AGP. Microsomal proteins from SPO-AGP-expressing cells were subjected to the second-round two-phase partition assay in the absence or presence of PI-PLC. Both SPO-AGP and V-PPase, which is an integral membrane protein, were detected by immunoblotting. Each lane in these figures contained proteins corresponding to an equal amount of microsomes or medium.

## Discussion

In this work we investigated the role of the GPI anchor in the transport and modification of NtAGP1, a classical AGP in tobacco BY-2 cells. The expression of the GFP fusion protein allowed us to detect the smear-migrating GFP fusion protein with low mobility on SDS-polyacrylamide gels (Fig. 1). This behavior of the fusion protein was consistent with the typical migration pattern of classical AGP in many plant species (e.g., Putoczki et al. 2007; Lind et al. 1994; Maurer et al. 2010). In contrast, the fusion protein formed using another fluorescent protein, mKikGR (Habuchi et al. 2008), yielded one slow-migrating and two fast-migrating smear bands (Suppl. Fig. S1). To determine which of the observed migration behaviors of these fusion proteins represents the true nature of NtAGP1, we characterized another fusion protein with sporamin and found that this fusion protein migrated as a slow-migration smear band on SDS-polyacrylamide gels (Fig. 4). We therefore concluded that the slow migration and formation of a smear band on SDS-polyacrylamide gels are characteristic of NtAGP1. These observations as well as the observation of multiple bands of GFP-fused *Arabidopsis* AGP4, which is a member of the classical AGP family that is transiently expressed in *Arabidopsis* seedlings (Bernat-Silvestre et al. 2020) also suggested that differences in the non-glycosylated region of AGP as well as the expression level may affect the degree of modification of classical AGPs.

Our analyses of the localisation of GFP-AGP and SPO-AGP indicated that these proteins are predominantly localized to the PM (Figs. 2, 3, 5), and that both proteins are attached to the PM as GPI-anchored forms (Fig. 7). In addition, some of these proteins were present in intracellular structures (Figs. 3, 5) and in the medium. These observations suggested that NtAGP1 is a GPI-anchored PM protein with a small pool in an uncharacterized intracellular structure, possibly related to an as-yet uncharacterized intermediate component of AGP synthesis (Poulsen et al. 2014). Some other population of NtAGP1 is also released to the extracellular space following the cleavage of its GPI anchor, possibly via the actions of phospholipase in the extracellular space (Fig. 9, upper). Similar low-level secretion of AGP or fluorescence protein-tagged AGP was reported in *Arabidopsis* (Darjania et al. 2002; Bernat-Silvestre et al. 2021; Xue et al. 2017). Such a low-level secretion of AGP might be the result of a slow release of this protein into the medium, since we observed that the cycloheximide treatment did not cause a significant decrease of cellular SPO-AGP within 24 hr (Suppl. Fig. S4). The presence of LM6 signal in both the cell wall and intracellular structures (Fig. 5) suggested that such a distribution is not specific to NtAGP1; rather it is common to other AGPs expressed in tobacco BY-2 cells, although we cannot rule out a possibility that significant proportion of the LM6 signals represent the presence of rhamnogalacturonan I.

In contrast to the fusion proteins with an intact AGP with a GPI-anchoring signal, we detected mutant fusion proteins without a GPI-anchoring signal (i.e., GFP-AGPΔC and SPO-AGPΔC) as multiple bands at different migration positions on SDS-polyacrylamide gels (Figs. 1, 4). The glycosylated forms of these proteins that migrated the most slowly and the most readily formed smear bands were predominantly recovered in the medium fraction (Figs. 1, 4). The fractionation analysis of cells expressing these proteins indicated that the intermediate forms were recovered partly in the membranous organelles in the cells and partly in the periplasmic space (Figs. 1, 4). The migration positions of these proteins on SDS-polyacrylamide gels were slower than those of the calculated size of non-glycosylated forms of these proteins, suggesting that these proteins are modified to some extent, probably with glycans. Purification of these proteins and analyses of their sugar compositions will clarify whether such modification takes place.

Our fractionation study of microsomes from SPO-AGPΔC-expressing cells indicated that the intermediate SPO-AGPΔC was localized in the ER with partial localisation in the Golgi apparatus (Fig. 6). The smallest form of this protein, which was similar in size to SPO, was recovered in the soluble fraction from the cells, which consisted of the cytoplasm and vacuolar sap. The smallest form of SPO-AGPΔC was also recovered in the vacuoplast fraction. These observations indicated that some of the SPO-AGPΔC protein was transported to the vacuoles, while most of the AGP part of these proteins was cleaved off or degraded in the vacuoles (Fig. 9, lower).

These observations indicated that GPI anchoring is not only essential for the proper and efficient transport of NtAGP1 to the plasma membrane or extracellular space but also required for the efficient glycosylation. Some of the inefficient-modified forms, i.e., the intermediate forms, were retained in the ER, and some of them could also be transported to the vacuoles for degradation (Fig. 9, lower). These observations are partly consistent with the case of a Citrin-tagged and GPI-anchoring signal-deleted fasciclin-like AGP mutant (Xue et al. 2017). In that case, most of the mutant protein was retained in the ER. In our present experiments, however, vacuolar localisation was observed. This difference in vacuolar localisation between the Xue et al. report and our present study might be attributable to either the different detection systems or different proteins used.

It has been reported that the vacuoles are scavenger organelles that degrade damaged proteins and organelles. In yeast, it has been shown that vacuolar targeting is a mechanism for degrading improperly folded proteins in the ER, and this transport is mediated by a receptor for vacuolar targeting (Hong et al. 1996). In plants, targeting of soluble proteins to the lytic vacuoles (such as the central vacuoles in tobacco BY-2 cells) is mediated by VSR proteins (reviewed in Shimada et al. 2018). Large and hydrophobic residues, such as Ile and Leu, in the vacuolar-sorting determinants in proteins play a pivotal role in the targeting (Ahmed et al. 2000; Brown et al. 2003; Matsuoka and Nakamura 1995, 1999; Paris et al. 1997). It is thus interesting that inefficient glycosylation of SPO-AGPΔC may cause the exposure of a large and hydrophobic side chain in the mature region of AGP, and this characteristic of SPO-AGPΔC may have allowed the targeting of this protein to the vacuole after recognition by VSR proteins, as in the case of yeast (Hong et al. 1986). In fact, the SPLA sequence at aa positions 67–70 of the NtAGP1 precursor resembles the NPIR vacuolar sorting determinants of sporamin and aleurain propeptides (Matsuoka and Nakamura 1995, 1999; Paris et al. 1997), and this region may be a good candidate for a region that is exposed under inefficient glycosylation.

Not only the mislocalisation but also alteration of the glycan moiety was observed in the secreted SPO-AGPΔC from SPO-AGP (Fig. 8). Secreted SPO-AGPΔC was recognized by the mAB LM2, which was shown to recognize β-GlcA residues in AG glycan (Yates et al. 1996). It has been shown that the structure of the glycan moiety of artificial AGP secreted from tobacco BY-2 cells contains β-GlcA, which is further modified partly by rhamnose (Tan et al. 2004, 2010). In addition, the β-GlcA moiety of AGP is sometimes methylated by the action of DUF576-family methyltransferases (Temple et al. 2019). It is thus possible that (*i*) the glycan structure of secreted SPO-AGPΔC is similar to that of the artificial AGPs, which contain a terminal β-GlcA, whereas such a structure is absent in SPO-AGP, and (*ii*) this difference is achieved by an efficient modification with rhamnose or methyl residues. Further analyses of the sugar compositions of the purified SPO-AGP and secreted SPO-AGPΔC proteins are necessary to fully explore these possibilities.

**Fig. 8.**
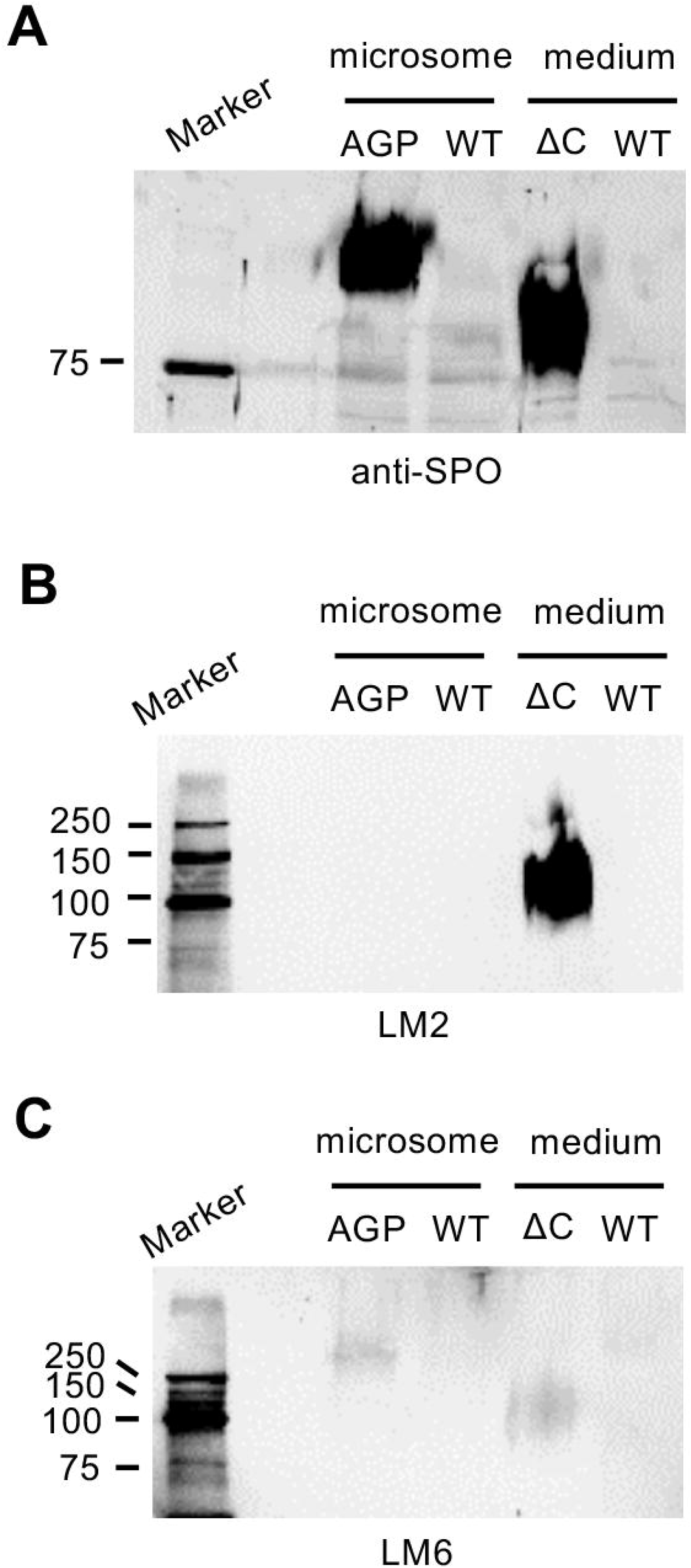
The large form of SPO-AGPΔC was specifically recognized by LM2 monoclonal antibody. **A:** The detection by anti-sporamin antibody showed that nearly equal amounts of sporamin-fusion proteins were loaded. **B:** Detection by the monoclonal antibody LM2, which recognizes a glycan epitope with b-linked glucuronic acid. **C:** Detection by the monoclonal antibody LM6, which recognizes a glycan epitope consisting of (1-5)-α-L-arabinosyl residues. AGP: microsomes from SPO-AGP-expressing cells, ΔC: culture medium from SPO-AGPΔC expressing culture, WT: corresponding microsomes and medium from non-transformed BY-2 cells with an equal amount of protein with either microsomes from SPO-AGP or culture medium from SPO-AGPΔC.

What causes the difference in glycan structure between SPO-AGP and secreted SPO-AGPΔC? One possible answer may be related to the difference in transport routes of these proteins in the late secretory pathway, with different distributions of the enzymes responsible for modification for the AGP glycan. A study of tobacco protoplasts using a GFP-tagged pectin methylesterase inhibitor protein and its mutant without a GPI-anchoring signal revealed that the late transport pathways of these proteins are different (De Caroli et al. 2011). The DUF576-family methyltransferases that act against AGP glycan have been detected not only in the Golgi apparatus but also in small punctate structures that are distinct from the Golgi apparatus (Temple et al. 2019). Similarly, a subset of glycosyltransferases involved in AGP glycan biogenesis was identified in a non-Golgi and non-TGN (trans-Golgi network) punctate structure (Poulsen et al. 2015). An analysis of the secretome of *Arabidopsis* indicated that there are at least two distinct secretory pathways in *Arabidopsis* leaf cells (Uemura et al. 2019). One of these pathways, which depends on SYP4-type SNAREs, is involved in the secretion of many hydrolases (Uemura et al. 2019). In an analysis focusing on AGP in the data generated and described in that paper (PRIDE dataset identifier PXD009099 by H. Nakagami et al.), only a small fraction of AGPs with a GPI-anchoring signal (18%) showed significantly high abundance in apoplasts in a wild-type relative to a mutant with SYP4-type SNARE proteins (Suppl. Table S3).

In contrast, when the same approach was taken against two groups of secretory hydrolases, i.e., pectic-lyase like proteins without a GPI-anchoring signal and β-galactosidase, a greater proportion of these proteins showed a high level of secretion in the wild-type compared to the mutant SYP4-type SNARE proteins (35% and 38%, respectively). In *Arabidopsis*, the SYP4-type SNAREs are localized in not only the TGN adjusting the Golgi apparatus but also a small punctate structure that is called the Golgi-independent TGN or free TGN (Viotti et al. 2010; Kang et al. 2011; Uemura et al. 2014). A similar structure called a secretory vesicle cluster (SVC), which is involved in pectin secretion, is present in tobacco BY-2 cells (Toyooka et al. 2009). Analyses of the proteome and glycome of secretory vesicles marked with SYP61 indicated that these secretory vesicles are rich in the SYP4-type SNARE proteins as well as glycans that are recognized by mABs against pectin (Drakakaki et al. 2012; Wilkop et al. 2019). Although glycans in the same vesicle fraction are also recognized by mABs against AGP, this recognition is weaker than that in the cell wall fraction, in contrast to the recognition by mABs that recognize pectin (Fig. 3 in Wilkop et al. 2019).

Collectively, these observations suggest that the GPI-anchored AGP precursor would be predominantly transported from the TGN to an uncharacterized punctate organelle, where a subset of glycosyltransferases and/or methyltransferases localize (Poulsen et al. 2015), and would then be transported to the PM after efficient modification of the glycan in the compartment (Fig. 9 AGP). The absence of a GPI anchor in the AGP precursor prevents taking this route from the TGN, and it passes through the default secretion pathway; this mis-sorting limits the final maturation of AGP glycan.

**Fig. 9.**
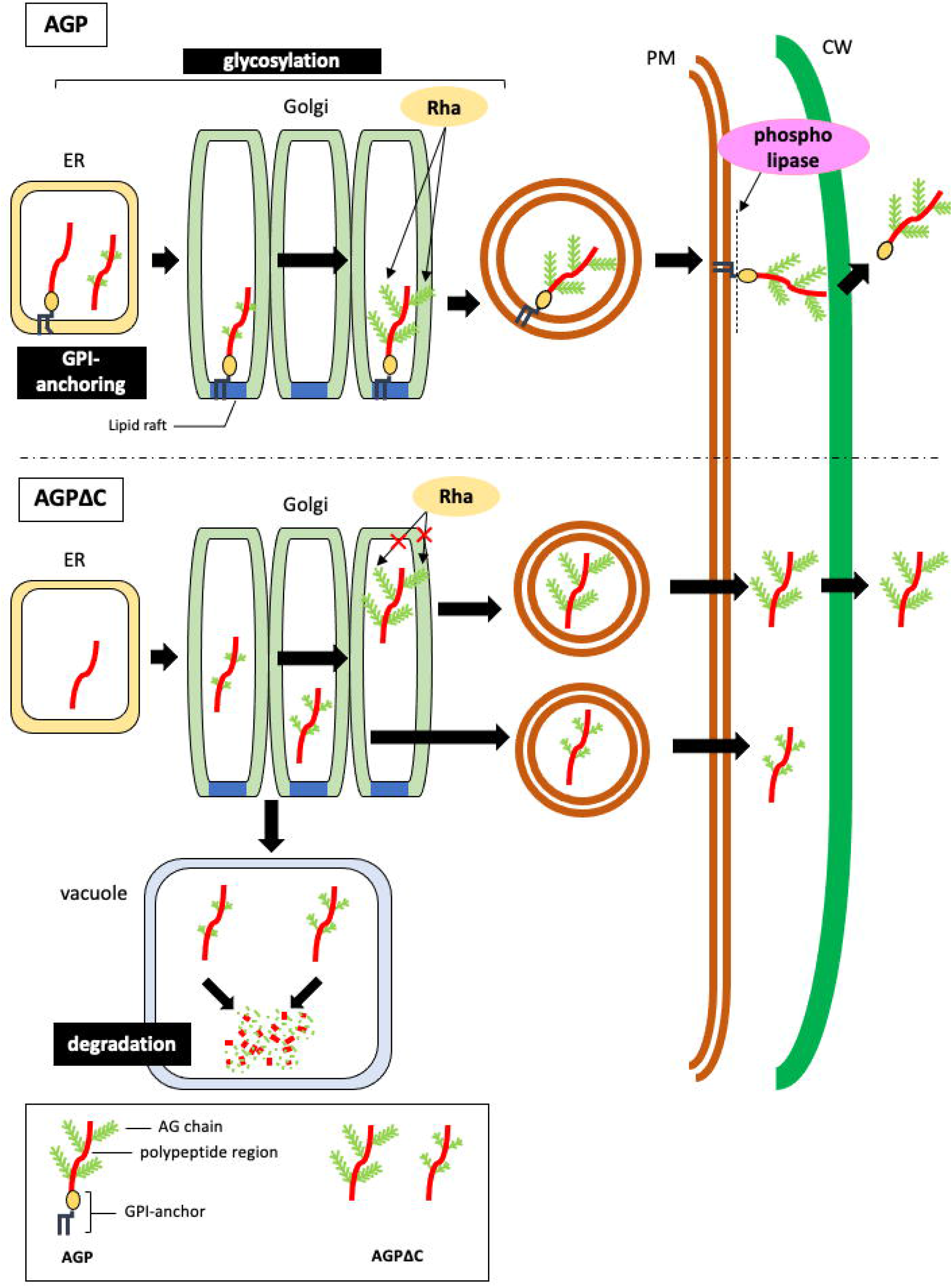
Proposed model of the transport and localisation of native NtAGP1 and its mutant lacking the GPI-anchoring signal.

Another possible explanation for the difference in glycan structure is that the accessibility to the modification enzymes might not be efficient in the absence of a GPI anchor. Because most of the glycosyltransferases and methyltransferases in the secretory pathway are integral membrane proteins whose catalytic domains are located proximately to the phospholipid bilayer, the GPI anchoring of the AGP precursor will bring the protein close to the catalytic sites of these enzymes. Without a GPI anchor, such access would not be efficient, and this characteristic would prevent the efficient maturation of the glycan side chain. A third possibility is that the transport speed is different between SPO-AGP and SPO-AGPΔC, and this difference contributes to the difference in the glycan side chain. Future studies to address the pathways and kinetics of transport and modification of AGP using SPO-AGP as a model will reveal which scenario or combination of scenarios is most likely.

## Materials and Methods

An outline of the methods and critical information about the materials used in this study are provided next; more detailed information about the plasmid construction, cell fixation, cell fractionation, antibodies and immunological methods, microscopy, two-phase separation, and other assays are described in the Supplemental Materials and Methods.

### Construction of plasmids

A tobacco cDNA clone, BY28237 (GenBank accession no. LC128049.1) encoding the NtAGP1 protein (GenBank accession no. BAU61512) was used for the construction of plasmids used herein. The fusion protein of GFP and NtAGP1, designated as GFP-AGP, was designed using this clone and a mutant GFP, i.e., sGFP(S65T) (Chiu et al. 1996). For the construction of sporamin fusions, the coding sequence for the signal peptide and the mature sporamin fusion construct (Δpro sporamin; Matsuoka and Nakamura 1991) was used for the signal peptide-sporamin region. All of the expression constructs were subcloned into plant expression vector pMAT137 (Yuasa et al. 2005) and used for transformation. The detailed methods used for the construction of plasmids are described in the Supplemental Materials and Methods.

### Culture, transformation, fixation, and immunostaining of tobacco BY-2 cells

The cultures and transformation of tobacco BY-2 cells and their transformants were maintained as described (Tasaki et al. 2014). For immunostaining, BY-2 cells or its transformants were fixed and permeabilized and incubated with primary and secondary antibodies essentially as described (Yuasa et al. 2005). Seven-day-old cells (stationary phase) were used in all of the experiments except when stated otherwise in the figure legends.

### Confocal microscopy and image analysis

The localisations of GFP-AGP and GFP-AGPΔC expressed in transformed BY-2 cells were visualized using a confocal laser scanning microscope (TCS SP8, Leica, Wetzlar, Germany) essentially as described (Tasaki et al. 2014). In some cases, Z-stack images were collected and converted to a rotating movie. To visualize the plasma membrane using FM 4-64 or cytoplasm using chloromethyl SNARF-1, cells were stained with either FM4-64 or chloromethyl SNARF-1 acetate as described (Yuasa et al. 2005). Confocal images of the red fluorescence of stained cells were collected as described above with adjustment of the excitation and emission wavelengths.

### Antibodies

Rabbit anti-mitochondrial porin antibserum was generated as described in the Supplemental Materials and Methods. Antisera against SDS-denatured SPO (Matsuoka and Nakamura., 1991), affinity purified anti-native SPO (Matsuoka et al. 1995), affinity purified anti-plant Sec61 (Yuasa et al. 2005), anti-V-PPase (Toyooka et al. 2009), anti-PIP (Suga et al. 2001), and anti-GLMT1 (Liu et al. 2015) antibodies were used at the appropriate dilutions. The monoclonal anti-AGP glycan antibodies LM2 and LM6 (Yates et al. 1996), were purchased from PlantProbes (Leeds, UK). For the visualisation of the antigen-antibody complex by immunoblotting or indirect immunostaining, Alexa Fluor 488 goat anti-rabbit IgG (H+L), Alexa Fluor 568 goat anti-rabbit IgG (H+L), Alexa Fluor 568 goat anti-mouse IgG (H+L), Alexa Fluor 568 goat anti-rat IgG (H+L), Alexa Fluor 647 goat anti-rat IgG (H+L), and Alexa Fluor 647 goat anti-rat IgM (μ chain) were purchased from Invitrogen (Eugene, OR, USA).

### SDS-PAGE and detection of fusion proteins

For the estimation of the size and extent of modification of the fluorescent fusion proteins, the proteins were separated by SDS-PAGE, and their fluorescence was recorded by direct scanning of the gel using an image analyser (Typhoon 9600, GE Healthcare Bio-Science, Chicago, IL) as described (Tasaki et al. 2014). For the detection of SPO fusion proteins and AGP glycan, immunoblotting was carried out as described (Oda et al. 2020) using appropriate primary and fluorescence-tagged secondary antibodies.

For the comparison of the reactivities of SPO-AGP and secreted SPO-AGPΔC against anti-glycan antibodies, we performed an adjustment of the loading of SPO-AGP in the microsomes and secreted large form of SPO-AGPΔC after semi-quantitative immunoblotting using anti-sporamin antibody. Thereafter the volumes of the microsomes and the medium that gave nearly identical intensity were used for comparison by immunoblotting using anti-glycan antibodies.

### Immunoprecipitation

Immobilized anti-native sporamin was prepared as described (Matsuoka et al. 1995). Immunoprecipitation was performed essentially as described by Matsuoka and Nakamura (1991), except that the immobilized antibody was used instead of the anti-sporamin serum and protein A Sepharose.

### Subcellular fractionation of cells

Cells and media were separated from the suspension culture by filtration. For the differential centrifugation analyses, cells were mixed with buffer and disrupted using a Parr Cell Disruption Bomb (Parr Instruments, Moline, IL). The disrupted suspension was subjected to differential centrifugation. The precipitates after the 1,000*g*, 10,000*g*, and 100,000*g* centrifugations as well as the supernatant of the final spin were designated as P1, P10, P100, and S fractions, respectively. For the fractionation of endomembrane organelles and the plasma membrane, microsomes prepared from cells were separated by linear sucrose density gradient centrifugation essentially as described by Matsuoka et al. (1997). Protoplasts were prepared as described by Nagata et al. (1981). Vacuoplasts and cytoplasts were prepared from protoplasts as described by Sonobe (1990).

### Fractionation of GPI-anchored proteins by two-phase separation using Triton X-114

Precondensation of the nonionic detergent Triton X-114 and the fractionation of GPI-anchored proteins were performed as described by Murata et al. (2012) with minor modifications. In some cases, 6.5 m units of PI-PLC (phospholipase C, phosphatidylinositol-specific from *Bacillus cereus*) (Sigma-Aldrich, St. Louis, MO) were included during the incubation.

### Cycloheximide treatment

Twenty mL of the 7-day old SPO-AGP culture was transferred to a 100-mL Erlenmeyer flask, and 200 μL of 1 mM cycloheximide and was added to the flask. The resulting flask was shaken in the dark at 26°C at 130 rpm. At 0, 3, and 24 hr of incubation, 1 mL of the culture was collected from the flask. Cells were collected by centrifugation, mixed with an equal amount of SDS-sample buffer, and lysed by sonication. After heating, equal volumes of protein-solubilized solution were subjected to SDS-PAGE and immunoblotting. As a control, water was used instead of cycloheximide solution.

### Enzymes and protein assays

The α-mannosidase activity was measured as described (Boller and Kende, 1979) using *p*-nitrophenyl phosphate as a substrate. Protein concentrations were measured using a DC protein assay kit (Bio-Rad, Hercules, CA).

## Accession number

NtAGP1 cDNA sequence: LC128049.

## Acknowledgments

We thank Dr. Koji Yuasa of the RIKEN Plant Science Center for the production of recombinant mitochondrial porin and Mr. Nweke A. Boniface for improving the manuscript.

## Author contributions

YT, RT, and KM performed the experiments on GFP fusion proteins. DN, YS, MF, and KM performed the experiments on sporamin-fusion proteins. DN, YT, YS, and KM designed the experiments and analysed the data. KM conceived the project and wrote the article with contributions from all of the authors.

