## Supplementary figures and images for "Glycosylphosphatidylinositol-anchoring is required for the proper transport and glycosylation of classical arabinogalactan protein precursor in tobacco BY-2 cells"

### Supplemental figure S1

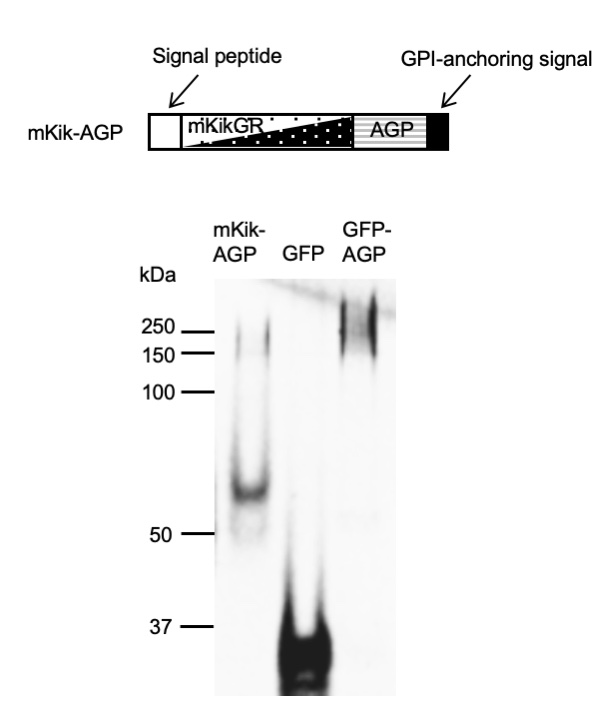

### Supplemental figure S2

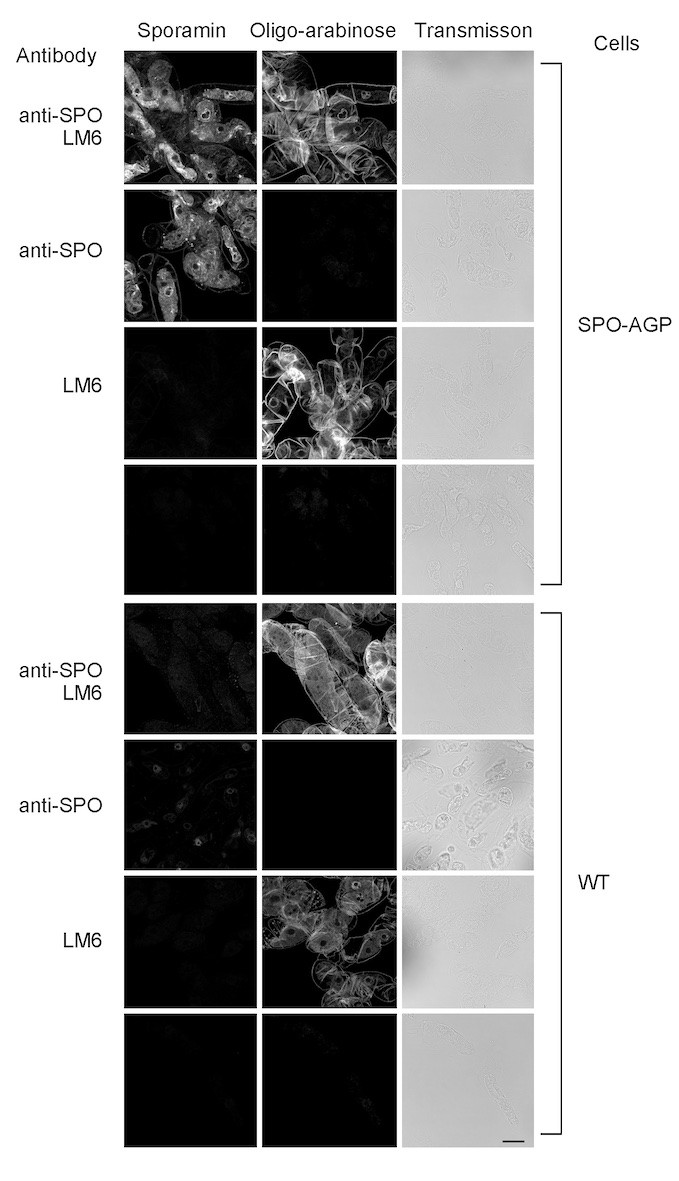

### Supplemental figure S3

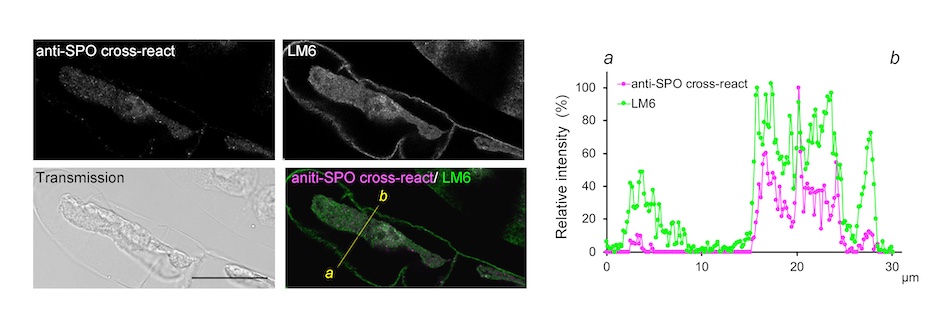

### Supplemental figure S4

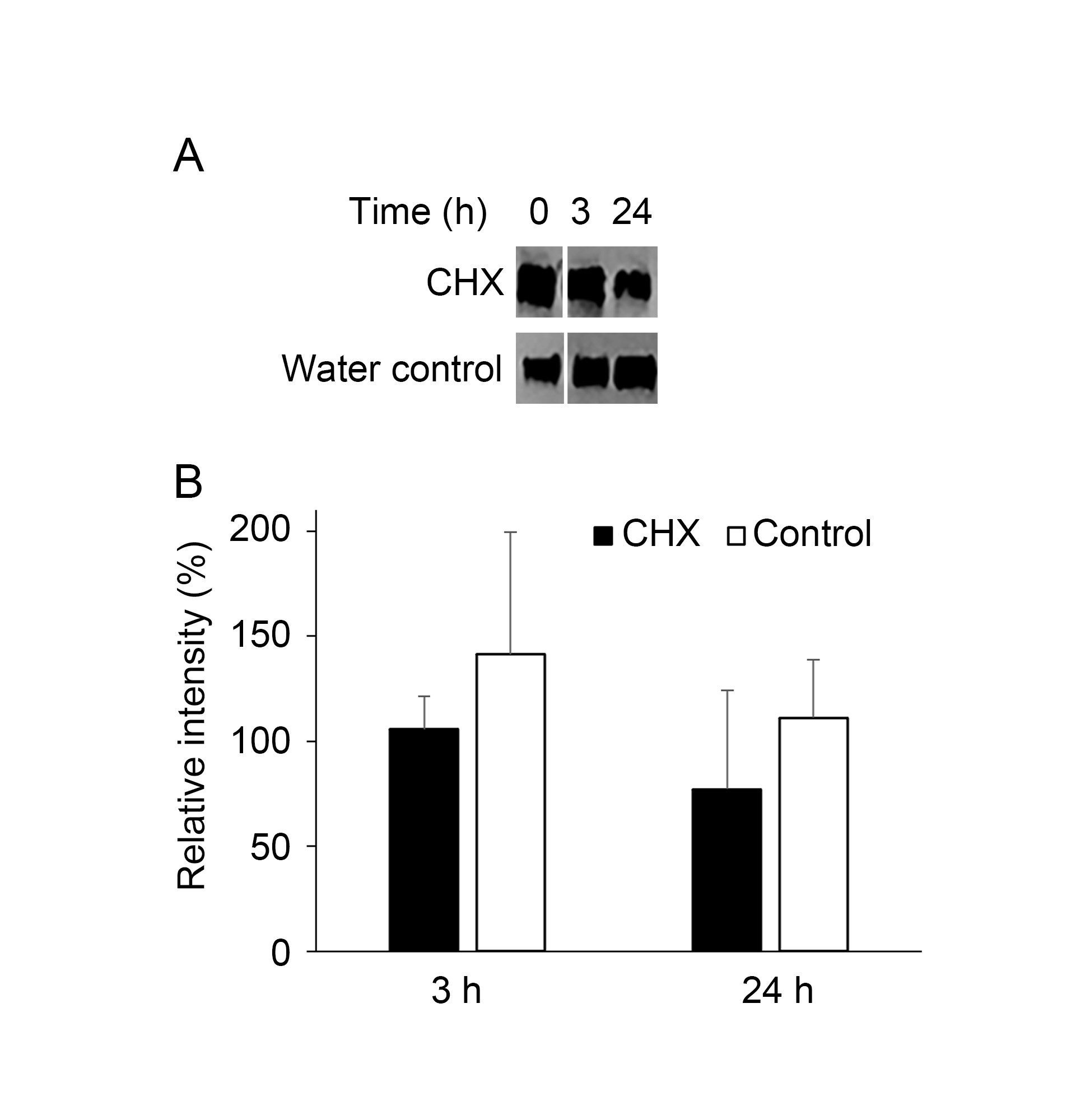
