## Supplementary material for "Glycosylphosphatidylinositol-anchoring is required for the proper transport and glycosylation of classical arabinogalactan protein precursor in tobacco BY-2 cells": Legends to Suppl Fig. and movie

**Legends to the Supplemental Figures and Movie**

**Suppl. Fig. S1.** Expression of the monomeric kikume green-red (mKikGR)-AGP fusion protein in tobacco BY-2 cells. Proteins were extracted from transformed tobacco BY-2 cells expressing mKikGR-AGP, GFP, or GFP-AGP and separated by SDS-PAGE. The green fluorescence in the gel emitted from these proteins was then recorded using a Typhoon 9600 image analyser.

**Suppl. Fig. S2.** Immunostaining of SPO-AGP-expressing cells (SPO-AGP) and non-transformed tobacco BY-2 cells (WT) with anti-sporamin and/or LM6 antibodies to detect sporamin-related epitopes and oligo-arabinose epitope, respectively. Confocal fluorescence images were collected using a Leica SP8 microscope with the same condition for all samples, and the adjustment of the intensities of the images was done with identical conditions for all images. All images are at the same magnification. Bar at the bottom right image: 30 mm.

**Suppl. Fig. S3.** Immunostaining of non-transformed tobacco BY-2 cells (WT) with anti-sporamin and/or LM6 antibodies to detect cross-react signals by anti-sporamin antibody and oligo-arabinose epitope, respectively. Confocal fluorescence images were collected using a Leica SP8 microscope with the same conditions for all samples, and the adjustment of the intensities of the images was done with identical conditions for all images. All images are at the same magnification. Bar at the bottom right image: 30 mm.

**Suppl. Fig. S4.** Seven-day culture of SPO-AGP-expressing cells was mixed with a 1/100 volume of cycloheximide (CHX) solution in water at the final concentration of 10 mM or in solvent control (water) and further cultured. At the indicated times, an aliquot of the culture was taken and cellular SPO-AGP was quantified after detection by western blotting. **A:** representative gel image. **B:** the results of the quantification of three independent experiments. Relative intensity to the zero time is shown. No significant difference was observed between CHX and water control by Student's t-test. Bars: SD.

**Suppl. Movie S1.** Three-dimensional construction of the distribution of GFP-AGP fluorescence in transformed tobacco BY-2 cells. Z-stack images of green fluorescence were collected and reconstituted to the 3D image. The constructed image was rotated.
