## Supplemental Materials and Methods for "Glycosylphosphatidylinositol-anchoring is required for the proper transport and glycosylation of classical arabinogalactan protein precursor in tobacco BY-2 cells"

**Materials and Methods**

*Construction of plasmid*

The pilot 5'-sequence of clones from the expressed sequence tag (EST) of tobacco BY-2 cells (Galis et al. 2006) allowed the identification of a clone (BY28237, GenBank accession no. LC128049.1) encoding the NtAGP1 protein (GenBank accession no. BAU61512), which is a typical AGP precursor that is nearly identical to a *Nicotiana alata* AGP precursor (Du et al. 1994). The signal peptide and GPI-anchoring signal of this protein were predicted using SignalP (http://www.cbs.dtu.dk/services/SignalP-3.0/) and PredGPI (http://gpcr.biocomp.unibo.it/predgpi/), respectively. The expression constructs used in this study were generated with the use of this information and the PCR primers listed in Supplemental Table S2.

The fusion protein of GFP and NtAGP1, designated as GFP-AGP, was designed as follows. The signal peptide and two amino acids from the cleavage site of the signal peptide were linked to a mutant GFP, i.e., sGFP(S65T) (Chiu et al. 1996), and then the C-terminus of this GFP was linked to the mature part and the GPI-anchoring signal of NtAGP1 by joining PCR. Using the EST as a template and primers P1 and P2, the coding region for the signal peptide and two amino acids with the N-terminal region of the GFP coding sequence was amplified. With the use of an sGFP(S65T)-containing plasmid as a template and primers P3 and P4, a DNA fragment encoding part of a signal peptide, two amino acids of mature NtAGP1, GFP, and the N-terminal part of mature NtAGP1 were amplified. Using the EST as a template and primers P5 and P6, a fragment encoding the C-terminal amino acids of GFP fused with the mature NtAGP1 and GPI-anchoring signal was amplified.

After these three fragments were mixed, the fusion construct was amplified using P1 and P6 primers. After the resulting fragment was digested with BglII and EcoRI restriction enzymes (because the primers also contain sequences for these enzymes), the fragment was cloned into the corresponding sites of plant expression vector pMAT137 (Yuasa et al. 2004). Using this plasmid as a template and primers P1 and P7, a fragment encoding GFP-AGPΔC was amplified. The resulting fragment was digested with BglII and EcoRI and cloned into pMAT137 as described above.

For the construction of sporamin fusions, the coding sequence for the signal peptide and mature sporamin fusion construct (Δpro sporamin; Matsuoka and Nakamura 1991) was used for the signal peptide-sporamin region. Using P8 and P9 as primers and a plasmid with the Δpro sporamin construct (pMAT108; Matsuoka and Nakamura, 1991) as a template, the region for the signal peptide-sporamin with the N-terminal region of mature NtAGP1 was amplified. By using the GFP-AGP-expressing plasmid described above as a template and primers P10 and P11, we amplified a fragment encoding the C-terminal region of sporamin with mature AGP, plus a GPI-anchoring signal. After these amplified fragments and primers P8 and P11 were mixed, a fragment encoding the SPO-AGP fusion protein was amplified. The resulting fragment was digested with BglII and EcoRI and cloned into pMAT137 as described above. Using this plasmid as a template and primers P8 and P12, a fragment encoding SPO-AGPΔC was amplified. The resulting fragment was digested with BglII and EcoRI and cloned into pMAT137 as described above.

*Culture and transformation of tobacco BY-2 cells*

The plant expression plasmids generated for the expression of AGP fusion proteins were used to transform tobacco BY-2 cells with *Agrobacterium* EHA 105 (Hood et al. 1993) as described (Tasaki et al. 2014). Cultures of tobacco BY-2 cells and their transformants were maintained as described (Matsuoka and Nakamura 1991). Seven-day-old cells (the stationary phase of growth) were used in all of the experiments except when stated otherwise in the figure legends.

*Confocal microscopy and image analysis*

The localisations of GFP-AGP and GFP-AGPΔC expressed in transformed BY-2 cells were visualized with a confocal laser scanning microscope (TCS SP8, Leica, Wetzlar, Germany). For the collection of fluorescence images from intact cells, the cell suspension was mounted on a slide glass attached to double-sided tape (Nicetack NW-40, Nichiban Co. Tokyo) with a 22-mm-dia. hole made by a paper punch. The cells were visualized using an HCPLAPOCS2 40×/1.30 OIL lens at a confocal pinhole setting of 65.3 μm. GFP fluorescence was excited by a 488-nm excitation laser beam of an Ar laser at 40% power, and the fluorescence between 530 and 575 nm was recorded using a hybrid detector at a setting of gain 406. Fluorescence of either FM 4-64 or 5- (and 6-) chloromethyl SNARF-1 was excited by a 514-nm excitation laser beam of an Ar laser, and the fluorescence between 650 and 700 nm was recorded using a hybrid detector at a setting of gain 100. For the capture of snapshot images at 1,024 × 1,024 pixels, we used the average of four scans using the linear scan mode at 100 Hz. For the collection of Z-stack images, the step size was optimized by Leica Application Suite software, and the scans were carried out as single scans using the linear scan mode at 100 Hz. For multi-colour imaging, the 'between lines' setting was used.

For the immuno-localisation analysis, fixed and immuno-stained cells were mounted as described above and visualized using an HCPLAPOCS2 63×/1.40 OIL lens at a confocal pinhole setting of 95.6 μm. Images at 1,024 × 1,024 pixels were collected as an average of four scans using the linear scan mode at 100 Hz without zooming or with zoom factor 2. Alexa Fluor 488 was excited at 488 nm using a white light laser at 60% power, and the fluorescence between 530 and 560 nm was recorded using a hybrid detector at a setting of gain 250 with a gating start at 3 ns and gating end at 6.5 ns. Alexa Fluor 568 was excited at 568 nm using a white light laser at 30% power, and the fluorescence between 600 and 640 nm was recorded using a hybrid detector at a setting of gain 100 with the gating start at 1.5 ns and the gating end at 6 ns. In all image collection cases, differential interference contrast (DIC) images at optimum exposure were collected with fluorescence images using a photomultiplier tube (PMT) detector. For multi-colour imaging, the 'between lines' setting was used.

For the reconstitution of the three-dimensional images from Z-stack images, the 3D Viewer software in the Leica Application Suite was used. After the adjustment of contrast and setting rotation, the movie was exported from this software. For the generation of composite images from Z-stack images, Z-project in the Fiji package of Image J2 (<https://imagej.net/software/fiji/>) was used. For the quantification of intensities on lines, a line nearly along the short axis of each cell clump was drawn by hand on the multi-colour image opened by Fiji, and the gray value of each colour along this line was collected. To compare the distribution of two different colours, the absolute values of intensities were converted to relative intensities with the highest intensity being 100%.

*Antiserum for mitochondrial porin*

A recombinant protein corresponding to the 14th to 165th positions of tobacco mitochondrial porin, encoded by the tobacco EST clone BY5432 (GenBank BP133193; Matsuoka et al. 2004) was expressed as a T7- and His_6_- tagged protein in *E. coli* after subcloning the corresponding fragment into a pET23 expression vector. The recombinant protein was purified using the His_6_-tag and used as an antigen to immunize a Japanese white rabbit. Antiserum from this rabbit was then used as anti-mitochondrial porin.

*Subcellular fractionation by differential centrifugation*

Cells and media were separated from the suspension culture by filtration. A 10-g aliquot of the cells was mixed with 10 mL of buffer containing 0.45 M sucrose, 50 mM Tris-MES (pH 7.3), 2 mM DTT, 1(w/v)% Na-ascorbate, and a piece of a cOmplete™ Mini EDTA-free tablet (Roche, Basel, Switzerland), and then disrupted by high-pressure N_2_ using a Parr Cell Disruption Bomb (model# 4636, Parr Instruments, Moline, IL) under 500 psi for 20 min. The disrupted suspension was centrifuged at 1,000*g* for 10 min at 4°C, and the supernatant was used as a total cellular fraction. For the differential centrifugation study, the supernatant was centrifuged at 10,000*g* for 10 min at 4°C.

Next, the supernatant was centrifuged at 100,000*g* for 1 hr at 4°C, then collected and supplemented with 1×TBS up to 1 mL as a soluble fraction. The precipitates after the 1,000*g*, 10,000*g*, and 100,000*g* centrifugations were resuspended with an equal volume of sonication buffer containing 1×TBS (25 mM Tris-HCl pH 7.4, 0.15 M NaCl), 1 mM EDTA, 1% Na-ascorbate, 100 µM leupeptin, 1 mM PMSF/0.5% isopropanol, and 1 mM NEM and sonicated using a Bioruptor (Cosmo Bio, Tokyo); the sonication condition consisted of power M at intervals of 0.5 min for 8 min. These sonicated suspensions were filled with 1×TBS to a final volume of 1 mL and used as the P1, P10, and P100 fractions, respectively. These fractions were used for immunoprecipitation and analysed by immunoblotting.

*Preparation of microsomal fractions and fractionation using sucrose density gradient centrifugation*

The preparation of microsomal fractions and the fractionation of microsomes using linear sucrose density gradient centrifugation was performed essentially as described by Matsuoka et al. (1997), except that a Beckman SW28.1 rotor was used. The continuous gradient was centrifuged at 100,000*g* for 18 hr at 4°C. After centrifugation, 1-mL fractions were collected from the bottom to the top of the gradient.

*Fractionation of GPI-anchored proteins by two-phase separation using Triton X-114*

The precondensation of Triton X-114 and the fractionation of GPI-anchored proteins were performed as described by Murata et al. (2012) with the following modifications. First, 100 μL of the microsomal fraction was gently mixed with 380 μL of 1×TBS and 120 μL of ice-chilled precondensed Triton X-114, and then incubated at 4°C overnight. The sample was centrifuged at 17,360*g* for 25 min at 4°C to remove detergent-insoluble materials. The supernatant was incubated at 37°C for 1.5 hr and centrifuged at 13,000*g* for 5 min at room temperature to induce phase separation. The upper phase (aqueous phase) and the lower phase were collected. The lower phase was gently mixed with 400 μL of chilled 1×TBS, and then the phase separation procedure was repeated and the upper phase in this second-round separation was combined with the first upper phase and used as an aqueous phase fraction. The resulting detergent-rich phase was mixed with a 4-fold volume of ice-cold acetone and incubated at −20°C overnight.

The proteins were recovered by centrifugation at 13,000*g* for 30 min at 4°C, and the precipitate was collected and air-dried on ice. The precipitate was resuspended with 400 μL of 25 mM Tris-HCl (pH 7.4) and used as the det fraction. In some cases, 6.5 m units of PI-PLC (phospholipase C, phosphatidylinositol-specific from *Bacillus cereus*) (Sigma-Aldrich, St. Louis, MO) were included during the incubation. For the processing of SPO-AGP, 200 μL of the det fraction was incubated with 1.8 m units of phospholipase C at 37°C for 2 hr, and thereafter the two-phase separation with Triton X-114 was conducted.
